# Ketamine modulates a norepinephrine-astroglial circuit to persistently suppress futility-induced passivity

**DOI:** 10.1101/2022.12.29.522099

**Authors:** Marc Duque, Alex B. Chen, Eric Hsu, Sujatha Narayan, Altyn Rymbek, Shahinoor Begum, Gesine Saher, Adam E. Cohen, David E. Olson, David A. Prober, Dwight E. Bergles, Mark C. Fishman, Florian Engert, Misha B. Ahrens

**Affiliations:** Department of Molecular and Cellular Biology, Harvard University; Cambridge, MA 02138, USA; Graduate Program in Neuroscience, Harvard Medical School; Boston, MA 02115, USA; Janelia Research Campus, Howard Hughes Medical Institute; Ashburn, VA 20147, USA; Solomon H. Snyder Department of Neuroscience, Johns Hopkins University School of Medicine, Baltimore, MD 21205, USA; Tianqiao and Chrissy Chen Institute for Neuroscience, Division of Biology and Biological Engineering, California Institute of Technology, Pasadena, CA, 91125, USA; Department of Physics, Harvard University; Cambridge, MA 02138, USA; Department of Chemistry and Chemical Biology, Harvard University; Cambridge, MA 02138, USA; Department of Neurogenetics, Max Planck Institute for Multidisciplinary Sciences, Göttingen 37075, Germany; Department of Chemistry, University of California, Davis; Davis, CA 95616, USA; Department of Biochemistry & Molecular Medicine, School of Medicine, University of California, Davis; Sacramento, CA 95817, USA; Center for Neuroscience, University of California, Davis; Davis, CA 95618, USA; Institute for Psychedelics and Neurotherapeutics, University of California, Davis, Davis, CA 95616, USA; Kavli Neuroscience Discovery Institute, Johns Hopkins University, Baltimore, MD 21205, USA; Department of Stem Cell and Regenerative Biology, Harvard University; Cambridge, MA 02138, USA

## Abstract

Mood-altering compounds hold promise for the treatment of many psychiatric disorders, such as depression, but connecting their molecular, circuit, and behavioral effects has been challenging. Here we find that, analogous to effects in rodent learned helplessness models, ketamine pre-exposure persistently suppresses futility-induced passivity in larval zebrafish. While antidepressants are thought to primarily act on neurons, brain-wide imaging in behaving zebrafish showed that ketamine elevates intracellular calcium in astroglia for many minutes, followed by persistent calcium downregulation post-washout. Calcium elevation depends on astroglial α1-adrenergic receptors and is required for suppression of passivity. Chemo-/optogenetic perturbations of noradrenergic neurons and astroglia demonstrate that the aftereffects of glial calcium elevation are sufficient to suppress passivity by inhibiting neuronal-astroglial integration of behavioral futility. Imaging in mouse cortex reveals that ketamine elevates astroglial calcium through conserved pathways, suggesting that ketamine exerts its behavioral effects by persistently modulating evolutionarily ancient neuromodulatory systems spanning neurons and astroglia.

## Main Text

Certain psychoactive compounds exert profound effects on behavioral state and mood that far outlast their lifetime in the body, providing a pharmacological means to treat diverse psychiatric diseases. In particular, ketamine (a dissociative NMDAR antagonist), MDMA (an empathogenic substrate of monoamine transporters), and psilocybin (a psychedelic) have been shown to reduce negative symptoms in several disorders, including major depressive disorder (MDD, or depression)^1–4^, obsessive compulsive disorder^5^, and post-traumatic stress disorder^6,7^. Despite the promise of long term adjustment of mood through brief pharmacological exposure, the mechanisms responsible for their behavioral effects remain poorly defined, and their widespread use has been limited by hallucinogenic side-effects and potential for abuse^8^. The pleiotropic effects of these drugs present a central challenge for further therapeutic development, as they act on receptors expressed on many different cell types that are widely distributed throughout the central nervous system (CNS). Furthermore, extrapolating from receptor engagement to behavior is extraordinarily challenging, due to our limited knowledge of brain circuitry and the many nonlinear processes associated with the generation and propagation of neural signals through these networks. Although the core psychological symptoms of psychiatric diseases like MDD can only be assessed in humans, animal models that incorporate key aspects of these syndromes provide the ability to perform mechanistic cellular manipulations to enable optimization of therapeutic action.

In the case of depression, a debilitating and widespread condition that is among the most prevalent causes of disability and places one of the largest burdens on public health worldwide^9,10^, the therapeutic landscape has changed dramatically with the discovery that ketamine and psilocybin have potent, fast-acting antidepressant effects. Ketamine and other fast-acting antidepressants rapidly ameliorate MDD symptoms in humans^1–4^ and, in rodents, rescue depression-related phenotypes, such as anhedonia and passive coping^11^. Functional measurements across species have characterized the effects of fast-acting antidepressants on neural activity in many brain areas, such as the prefrontal cortex^12^, habenula^13,14^, and serotonergic raphe^14^. These compounds exert complex responses in the CNS, influencing mTOR^15^, norepinephrine release and reuptake^16^, synaptic plasticity/metaplasticity associated with BDNF/TrkB signalling^17,18^, and extracellular matrix remodeling^19^, but the specific molecular targets and cellular populations responsible for the therapeutic effects of ketamine are unclear.

Recently, astroglia, a class of non-neuronal cells that play important roles in modulating neural activity^20–26^, has been implicated in the etiology of MDD^27,28^. Post-mortem anatomical analysis of MDD patient brains indicate that depression is associated with decreased astrocyte density in the hippocampus^29^ and cortex^30^, and in a corticosterone-induced rodent model of depression-like state, cortical astrocytes exhibited abnormal behavior-and serotonin-triggered responses^31^. Additionally, disruption to astroglial purinergic signaling contributes to depression-like phenotypes in rodents, and supplementing ATP chronically, or blocking acute ATP/adenosine signaling, leads to antidepressant-like effects^32–34^. However, models of antidepressant action involving astrocytes have not yet been developed.

Despite the available data on the effects of fast-acting antidepressants at the behavioral, circuit, and molecular levels, a mechanistic framework that bridges these different levels of analysis has remained elusive. While drug-induced changes in neuronal activity and synaptic plasticity can be readily defined in reduced systems or restricted areas of the CNS, it is difficult to extrapolate from this local focus to circuit and systems level changes more proximal to behavior.

The larval zebrafish has emerged as a vertebrate model system in which the brain-wide molecular and circuit effects of genetic and pharmacological perturbations can be linked to their impact on behavior. Their small size and optical transparency have enabled drug discovery^35,36^ and brain-wide imaging and perturbation of neural^37^ and glial^38,39^ activity at cellular resolution during behavior, enabling the generation of circuit hypotheses for behavior from sensory input to motor output. A subset of ketamine’s behavioral effects is conserved between zebrafish and mammals. Specifically, following CNS clearance, ketamine exhibits a sustained anxiolytic effect in adult zebrafish^40^ consistent with its effects in rodents. In juvenile zebrafish, ketamine inhibits passive coping in response to inescapable aversive shock^14^, a behavioral effect that manifests in rodents as a suppression of passivity in futility assays like the forced swimming and tail suspension tests^11^. However, despite these similarities in behavioral response to ketamine, as well as the high degree of conservation of receptors and underlying brain circuits between fish and mammals^41,42^, relatively few studies have been performed to examine the effects of ketamine and other fast-acting antidepressants on circuit dynamics at the whole-brain level. A previous study showed that inescapable stress recruits habenula firing and inhibits the serotonergic raphe to trigger passive coping and that ketamine inhibits habenular sensitivity to inescapable stress^14^, but neither the acute effect of ketamine on brain-wide neuronal activity nor its effects on non-neuronal cell types have been investigated. Another study used whole-brain light-sheet microscopy to show that antidepressants, including ketamine, affect functional neuronal connectivity in larval zebrafish^43^, but did not explore non-neuronal cells or the behavioral consequences of these changes.

An evolutionarily conserved behavior termed futility-induced switching, in which repeated failures to achieve goals or escape stressors lead to a persistent behavioral switch or a cessation of effort, can be interpreted as being due to discounting of effort’s expected value^44^. In humans, futility-induced behavioral changes can become maladaptive in MDD, often manifesting as a feeling of hopelessness^45^. In futility-related rodent assays, such as learned helplessness, the forced swim test, and the tail suspension test, behavioral outcomes are sensitive to both conventional and fast-acting antidepressant compounds^11,46^. Previous work has shown that larval zebrafish exhibit both passive coping and futility-induced passivity^14,38^. In futility-induced passivity, visual feedback from swim actions can be withheld so that swim attempts fail to trigger expected visual flow, as if animals fail to move through their environment despite neural motor commands being sent to the muscles, constituting an inescapable behavioral challenge. After tens of seconds of such motor futility, animals tend to ‘give up’ and become passive for similar durations. In this behavior, astroglia play central roles in integrating noradrenergic signals of swim failure and driving passivity^38^, raising the possibility that astroglial signaling might underlie some of the behavioral effects of fast-acting antidepressants.

Here, we leveraged the distinct advantages of larval zebrafish to delineate the effects of the fast-acting antidepressant ketamine on brain-wide neuronal activity and link these changes in cellular activity to behavior. We find that several pharmacological compounds, including ketamine, that are known to suppress futility-induced behavioral changes in mammals, also do so in larval zebrafish. We mapped the brain-wide physiological consequences of ketamine administration and found that it triggers strong, long-lasting, norepinephrine-dependent, astroglial calcium activity, followed by persistent calcium downregulation post-exposure, that is both necessary and sufficient to subsequently suppress futility-induced passivity *in vivo*. Our findings led us to predict that ketamine should similarly drive astroglial calcium activity in mammals through similar signaling pathways, a prediction we validated *in vivo* in rodents. Thus, ketamine acutely enhances calcium signaling in astroglia, the long-lasting consequences of which decrease the ability of behavioral futility to activate a norepinephrine-to-astroglia signaling axis that drives passivity, leading to a persistent brain state that promotes resilience.

### Ketamine pre-exposure persistently suppresses futility-induced passivity in larval zebrafish

We adapted a previously reported assay for futility-induced passivity in paralyzed, fictively behaving larval zebrafish^38^ to assess this behavior in unparalyzed, head-restrained fish (**Figure 1A**). Fish were embedded in agarose such that their heads were immobilized and their tails free to move. Using custom-built behavior apparatuses and software, we detected swimming in real time while delivering visual stimuli consisting of gratings that could move forward or backward relative to the fish. Trials (**Figure 1B**) consisted of three different time periods, ‘rest,’ ‘closed-loop,’ and ‘open-loop.’ During rest periods, gratings were stationary. During closed-loop periods, gratings steadily drift forward to induce swimming via the optomotor response, a position-stabilizing behavior^47,48^. If fish swim during closed-loop, the forward-moving gratings slow down or reverse direction, a visual feedback cue signaling forward self-motion; thus, swimming is perceived as being effective during closed-loop. In contrast, during open-loop trials, gratings drift forward with the same velocity as in closed-loop, but swimming does not change the velocity or direction of forward moving gratings. In this configuration, swimming is perceived as futile and constitutes an inescapable behavioral challenge similar to the rodent forced swim test. Open-loop induces a period of higher vigor swimming followed by transient cessation of swimming (passivity). The key behavioral variable assayed is the percentage of the open-loop period spent in the passive behavioral state. Furthermore, potential hyper-or hypo-locomotion-inducing effects of drugs are monitored through closed-loop passivity and closed-loop swim rate. We used this assay to test for persistent effects of psychoactive drugs on futility-related behavior.

**Figure 1.**
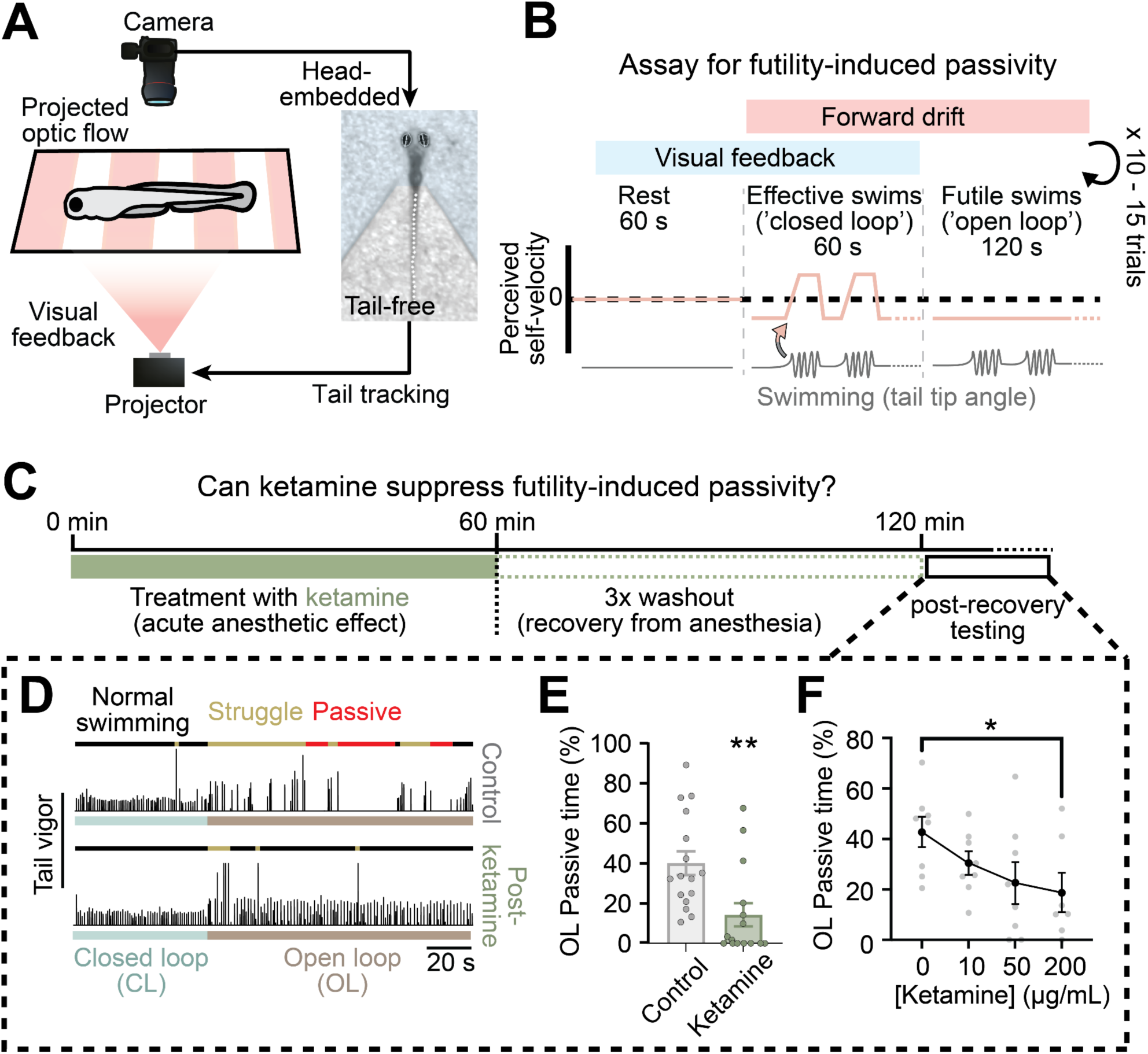
Ketamine suppresses futility-induced passivity in larval zebrafish. (A) Schematic of experimental workflow for imaging swimming behavior in unparalyzed larval zebrafish. Unparalyzed larval zebrafish are imaged and tail posture extracted in real time. Tail posture information is used to control a projector that displays optic flow stimuli (drifting gratings) to drive swimming. (B) Trial structure: 60 s of rest (no stimulus), 60 s of effective swimming (‘closed-loop’, forward optic flow, visual feedback), then 120 s of futile swimming (‘open-loop’, forward optic flow, no visual feedback), 10 to 15 repeated trials. (C) Timeline of experiments testing ketamine’s effect on futility-induced passivity. (D) Example trials for untreated control fish (top) and ketamine-treated fish (bottom). Colors above swim trace indicate normal swimming (black), struggle (yellow), and passivity (red). Colors below swim trace indicate closed-(teal) and open-loop (brown) periods. (E) Ketamine-treated fish spend less time passive (periods > 10 s with no swimming) than untreated controls during open loop. Mann Whitney test, 16 fish (control), 15 fish (ketamine), p = 0.0010. (F) Dose-response curve of ketamine’s suppression of open-loop passive time. Mann-Whitney test, N = 8 fish (control), 8 fish (ketamine), p = 0.0289. All error bars denote standard error of the mean (s.e.m.) * p < 0.05 ** p < 0.01.

Ketamine is an NMDA receptor (NMDAR) antagonist widely used as a rapid-acting dissociative anesthetic^49^. Previous studies have shown that at lower doses, after the dissociative effects disappear, ketamine elicits persistent antidepressant action that remain after washout of the drug^1,11^. To determine if futility-induced passivity behavior in zebrafish is altered by pre-treatment with ketamine, fish were exposed to 200 µg/mL ketamine (1 hour bath administration with a dose that decreases passive coping in response to inescapable behavioral challenge in older zebrafish^14^), then following washout allowed to recover from acute anesthetic effects. We then subjected these fish (and untreated clutch controls) to the futility-induced passivity assay (**Figure 1C**).

As a dissociative anesthetic, on acute timescales ketamine profoundly affects motor behavior, so we used two independent analyses to determine whether, after washout, ketamine had any lasting effects on baseline motor activity. First, we analyzed closed-loop swimming post-recovery and found that both treated and control fish exhibited normal swimming during closed-loop periods (**Figures 1D**, **S1A, S1B**; **Video S1**). Second, freely swimming fish exhibited identical swim rates between treated and untreated groups. Together, these data indicate that ketamine does not induce prolonged hyper-locomotion in larval zebrafish (**Figure S1C**). Having determined that ketamine does not alter baseline swimming, we analyzed its effects on futility-induced passivity in the open-loop period. During open loop, control fish exhibited periods of swimming and struggle characterized by larger, uncoordinated tail deflections (**Figures 1D** and **S1D**; **Video S1**), as well as periods of passivity characterized by a lack of swimming (**Figures 1D** and **1E**). Ketamine treatment significantly decreased the duration of passivity during open-loop (**Figures 1D and 1E**, **Video S2**) in a dose-dependent manner (**Figure 1F**). This decrease in passivity was not due to an inability of fish to swim or distinguish closed from open loop (**Figure S1E**), nor was it due to motor recovery from anesthesia, as the anesthetic sodium channel blocker MS-222 had no effect on futility-induced passivity (**Figures S1F-S1H**). Thus, ketamine suppresses futility-induced passivity in larval zebrafish in a manner distinct from its anesthetic or dissociative effects.

To assess the generalizability of these findings to other compounds, we characterized the pharmacological response profile of futility-induced passivity, using a panel of pharmacological agents known to affect, or to have no effect on, futility-induced behavior in other models (**Figures S1I-S1U**, **Table S1**). Compounds that have been previously reported to have fast-acting, antidepressants-like effects and that bind either NMDA receptors (ketamine, MK-801, DXM) (**Figures S1J and S1P**) or serotonergic receptors (DOI) (**Figure S1L**) caused a decrease in futility-induced passivity, although MK-801 also induced hyperlocomotion during closed loop (**Figure S1I**). The 5-HT2A receptor agonist, TCB-2^50^, which has not been characterized in rodent passivity models, also suppressed futility-induced passivity (**Figures S1I and S1L**), similar to the effects of other psychedelics (LSD, DOI and psilocybin^51,52^) in mammalian models. One of the drawbacks for therapeutic use of ketamine and classical psychedelics is their potential for hallucinogenic effects. This has led to the generation of psychedelic analogs with diverse chemical structures and pharmacological targets predicted to reduce these side effects^53–55^. We found that, as reported in rodent models, these compounds also significantly reduced passivity in our assay (**Figures S1Q**, and **S1R**), demonstrating a conservation of their behavioral effects across species. On the other hand, classical antidepressants fluoxetine or (S)-citalopram (SSRIs) and desipramine (TCAs) showed no passivity-suppressing effects after a single, one-hour dose, consistent with the multiple doses and prolonged onset time required for behavioral effects in rodents and humans^56,57^ (**Figure S1K**). However, (2R,6R)-hydroxynorketamine, a metabolite of ketamine that has been reported to decrease passivity in the rodent forced swim test^58^, had no effect in our futility-induced passivity assay (**Figure S1S**), raising the possibility that it acts through pathways not found in the larval zebrafish or has low bioavailability in this whole-animal assay. Additionally, we assayed other NMDAR antagonists, including memantine, a low-affinity NMDAR antagonist that binds to the same site as ketamine/MK-801, and AP5, a competitive NMDAR antagonist that binds to a distinct site. Neither of these compounds significantly altered futility-induced passivity at the doses tested (**Figures S1T and S1U**).

A single dose of ketamine and psychedelics can produce behavioral effects that far outlast their presence in the body. Therefore, we also assayed the persistent effects of drugs on passivity 24 hours after treatment and washout. Of the drugs tested, only the phencyclidine-site NMDAR antagonists ketamine and MK-801 retained a passivity-suppressing effect after 24 hours (**Figure S1M**), although animals treated with MK-801 maintained hyperlocomotion, complicating interpretation of MK-801 action. Classical antidepressants had no long-lasting effects on passivity (**Figure S1N**), as expected from their lack of acute single-dose efficacy, and while psychedelic 5-HT2AR agonists acutely suppressed passivity, this suppression was no longer present after 24 hours (**Figure S1O**). Therefore, ketamine triggers a persistent decrease in futility-induced passivity in the larval zebrafish, which could reflect lasting alterations in brain activity.

### Brain-wide effects of ketamine on neural activity and neuromodulation

The optical accessibility of the larval zebrafish offers unique opportunities for assessing how fast-acting antidepressants affect the activity of neurons and non-neuronal cells across the entire brain during behavior. We used light-sheet imaging to collect brain-wide neural activity profiles during acute ketamine exposure at single-cell spatial resolution in *Tg(elavl3:H2B-GCaMP7f)* fish (**Figure 2A**). Images were then registered to a zebrafish brain atlas^59^ to identify regions of concentrated activity (**Figures S2A and S2B**). Consistent with ketamine’s inhibitory effects on spontaneous and sensory-evoked locomotion, neural activity was inhibited in early visual areas, such as the optic tectum and hindbrain motor regions (**Figures 2B and 2C**). However, in a small subset of brain areas neuronal activity was enhanced by ketamine, including noradrenergic and serotonergic nuclei and several regions of the cerebellum (**Figures 2C-2E**). While ketamine’s effects on the cerebellum are relatively understudied, emerging evidence suggests that the cerebellum can play an important role in affect and affective disorders through its interconnectivity with cognitive, limbic, and monoaminergic nuclei^60,61^. Ketamine’s stimulatory effects on serotonergic and noradrenergic regions have been reported in mammals^16,62^, suggesting that the effects of ketamine on these evolutionarily ancient, monoaminergic nuclei are conserved in fish.

**Figure 2.**
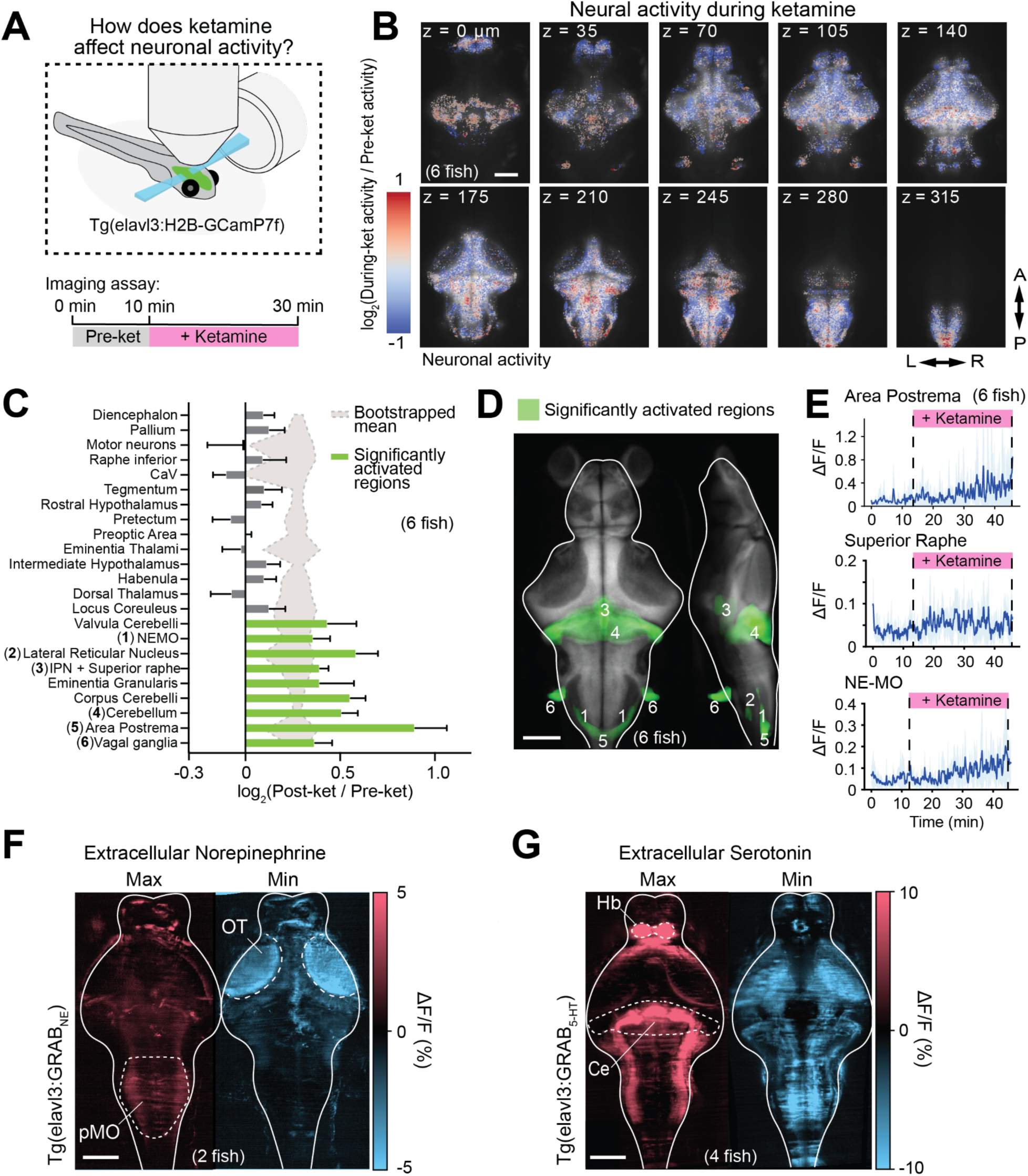
Activation of noradrenergic and serotoninergic populations following acute ketamine. (A) Fish expressing the nuclear calcium indicator H2B-GCaMP7f under a pan-neuronal promoter were imaged using a light-sheet microscope before and during acute treatment with ketamine. (B) Average activity changes across six registered fish (log-ratio of ΔF/F following ketamine administration to before administration) at different anatomical depths after ketamine treatment. (C) Z-Brain^59^ areas activated significantly above (green) or below (gray) chance (background shaded region, 95% CI) following ketamine treatment. Chance levels determined through bootstrapping with shuffle indices (nBoot = 10000). Numbers in parenthesis correspond to regions in (D). (D) Brain areas that were significantly activated following ketamine treatment in green overlaid onto the reference atlas fish. Labeled numbers correspond to regions listed in (C). (E) Average neural activity (ΔF/F) of indicated noradrenergic and serotonergic regions across fish (N=6 fish, top 10% cells for each region per fish). Shaded region denotes s.e.m. (F) Maximum and minimum z-projections for average NE levels across the brain during ketamine exposure imaged using GRAB_NE_ (N = 2 fish). OT = Optic Tectum, pMO = posterior Medulla Oblongata. (G) Maximum and minimum Z-projections for average 5-HT levels across the brain during ketamine exposure imaged using GRAB_5-HT_ (N = 4 fish). Ce = Cerebellum, Hb = Habenula. All scale bars 100μm.

As monoaminergic system dysfunction is highly correlated with depressive phenotypes, and monoaminergic systems are the primary targets of conventional pharmacological antidepressant treatments, we examined in more detail ketamine’s effects on the activity of noradrenergic and serotonergic nuclei in the brain. Noradrenergic neurons in a previously characterized medullary population (NE-MO, or MO-NE, likely area A2 neurons in mammals^63^) that drives futility-induced passivity, along with other noradrenergic clusters in the area postrema, all exhibited increases in activity during ketamine exposure (**Figures 2C-2E**). By contrast, activity in the locus coeruleus was suppressed (**Figure 2C**). In addition to the noradrenergic system, hindbrain serotonergic populations also increased their activity during acute ketamine exposure (**Figure 2E**), consistent with observations in rodents^64^. To complement the assessment of neural activity changes in monoaminergic nuclei, we also directly visualized changes in both norepinephrine (NE) and serotonin (5-HT) levels in the brain following ketamine treatment, by imaging pan-neuronally expressed extracellular sensors of NE^65^ (using the *Tg(elavl3:GRAB_NE_)* line) and 5-HT^66^ (using the *Tg(elavl3:GRAB_5-HT_)* line) (**Figures S2C** and **S2D**). Ketamine has been previously shown to elevate sympathetic drive and NE in the periphery^62,67^. We found that, consistent with its peripheral sympathomimetic effects, ketamine administration elevated NE across the hindbrain, especially in the posterior medulla oblongata (pMO) (**Figure 2F**, **Figure S2E**), while NE levels decreased in the optic tectum (OT) (**Figure 2F**, **Figure S2E**). Direct visualization of 5-HT revealed that ketamine induced strong elevation in the habenula (Hb) and cerebellum (Ce), but had more diverse effects in the optic tectum and medulla (**Figures 2G**, **Figure S2F**). Elevation of 5-HT in the habenula is particularly interesting, as serotonin has been implicated in the pathogenesis of depression-like behaviors and the fast-acting behavioral effects of ketamine in both rodents^13,68,69^ and fish^14^. Together, these data indicate that ketamine acutely alters monoaminergic neuromodulation across the brain in a region-dependent manner, raising the possibility that some of the effects of ketamine on futility-induced passivity may arise through noradrenergic or serotonergic signaling.

### Norepinephrine-dependent elevation of astroglial calcium by ketamine

Norepinephrine (NE) is known to be a major, evolutionarily conserved driver of astroglial calcium signaling^21–24,38^. Moreover, during futility-induced passivity in the larval zebrafish, hindbrain NE drives radial astroglial calcium, and astroglial calcium elevation suppresses futile swims^38^. To determine if ketamine acts on futility-related NE neuromodulation of astroglial calcium signaling, we imaged calcium activity in radial astrocytes during ketamine exposure using widefield fluorescence imaging in unparalyzed, head-embedded *Tg(gfap:jRGECO1a)* fish (**Figure 3A**). Strikingly, acute exposure to ketamine induced a strong prolonged rise in astroglial cytosolic calcium concentration (**Figures 3B-3D**; **Video S3**) that did not return to baseline for over 20 minutes (**Figures S3A and S3B**). The ketamine-induced calcium increase was many times larger and more prolonged than the transient calcium increases that occurred during open-loop struggles (magenta vs. black in **Figure 3D**). In contrast, fluoxetine, DOI, and MS-222 did not elevate cytosolic calcium in astroglia (**Figure 3E**).

**Figure 3.**
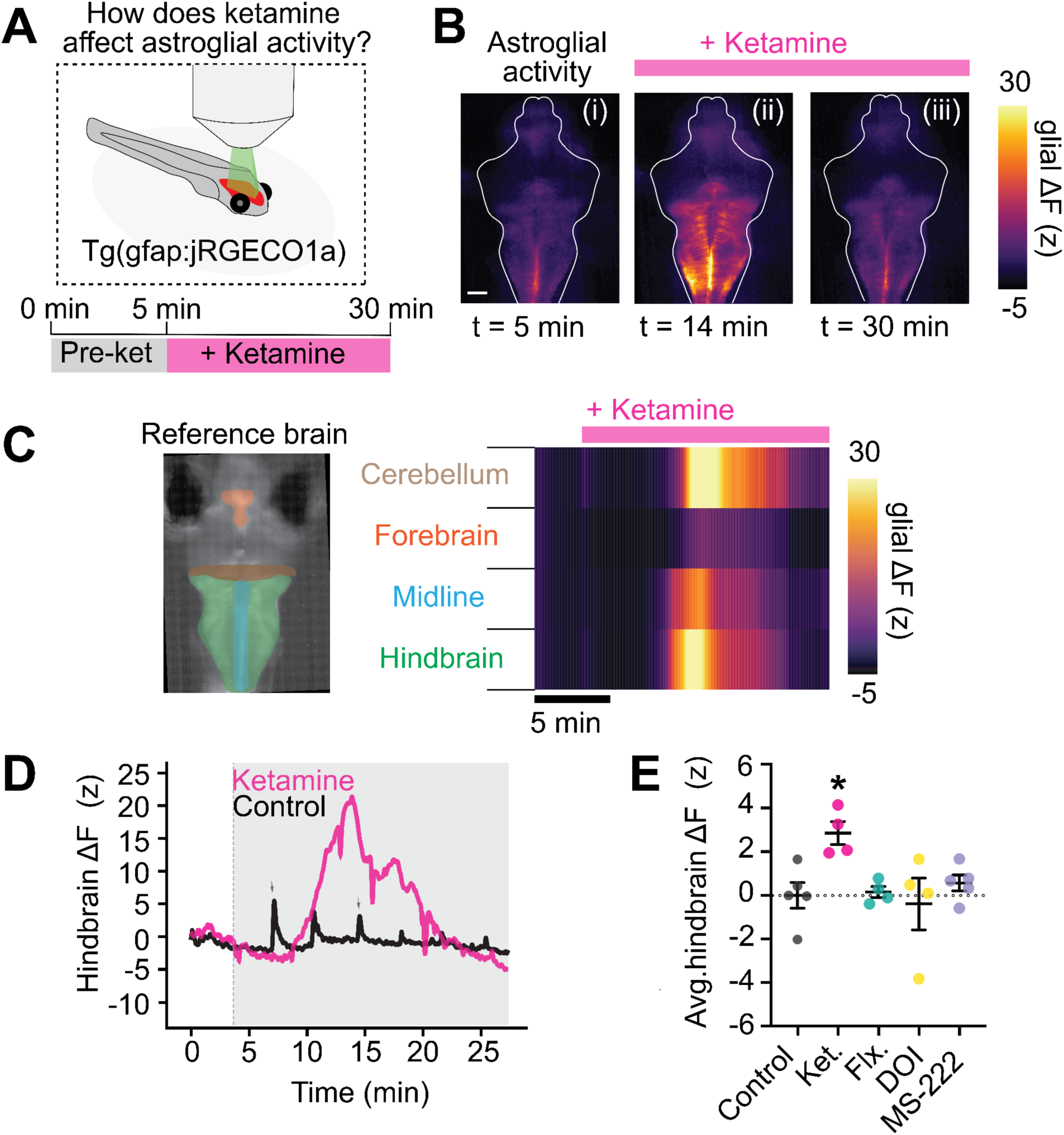
Ketamine triggers a long-lasting calcium elevation in astroglia. (A) Fish expressing the calcium indicator jRGECO1a in astroglia were imaged using an epifluorescence microscope before and during acute treatment with ketamine for 30 minutes. (B) Fluorescence micrographs of jRGECO1a signal in an example fish at three time points illustrating elevation and return to baseline of cytosolic calcium in presence of ketamine. Scale bar 50μm. (C) Heatmap of glial jRGECO1a signal in four ROIs (left) for fish in (C). Pink bar indicates ketamine in the bath. (D) Hindbrain jRGECO1a fluorescence change in an example fish treated with ketamine (200 μg/mL) or vehicle (control). Dips in magenta signal are imaging artifacts. Short increases in astrocytic calcium during struggles in control fish are indicated by gray arrowheads. (E) Average hindbrain fluorescence change following treatment with listed compounds. Ketamine, but not fluoxetine (SSRI), DOI (5-HT2AR agonist) or MS-222 (anesthetic) elevates cytosolic astroglial calcium. One-way ANOVA with Tukey’s multiple comparison test (vs. control). N = 5 fish (control, MS-222), 4 fish (all other conditions). p=0.0169 (Ketamine), 0.9395 (MS-222), 0.9893 (DOI), 0.9996 (Fluoxetine).

To determine whether the ketamine-induced astroglial calcium elevation depends on extracellular NE, we performed simultaneous two-color brain-wide imaging of intracellular calcium in astroglia as well as extracellular NE (**Figure 4A**) in fish expressing jRGECO1 in astroglia and GRAB_NE_ in neurons. NE elevation and astroglial calcium elevation had similar time courses (**Figure 4B**), and pharmacological inhibition of α1-adrenergic receptors (α1-ARs) (**Figure 4C**), Gq-coupled receptors primarily expressed by astroglia, with prazosin abolished the stimulatory effect of ketamine on astroglial calcium (**Figure 4D**). Moreover, pharmacological blockade of α1-AR signaling prevented the ability of ketamine to suppress futility-induced passivity (**Figure 4E**, **Figures S4A-S4C**). These data suggest that NE-dependent astroglial calcium regulation is a critical step in the reduction of futility induced passivity by ketamine.

**Figure 4.**
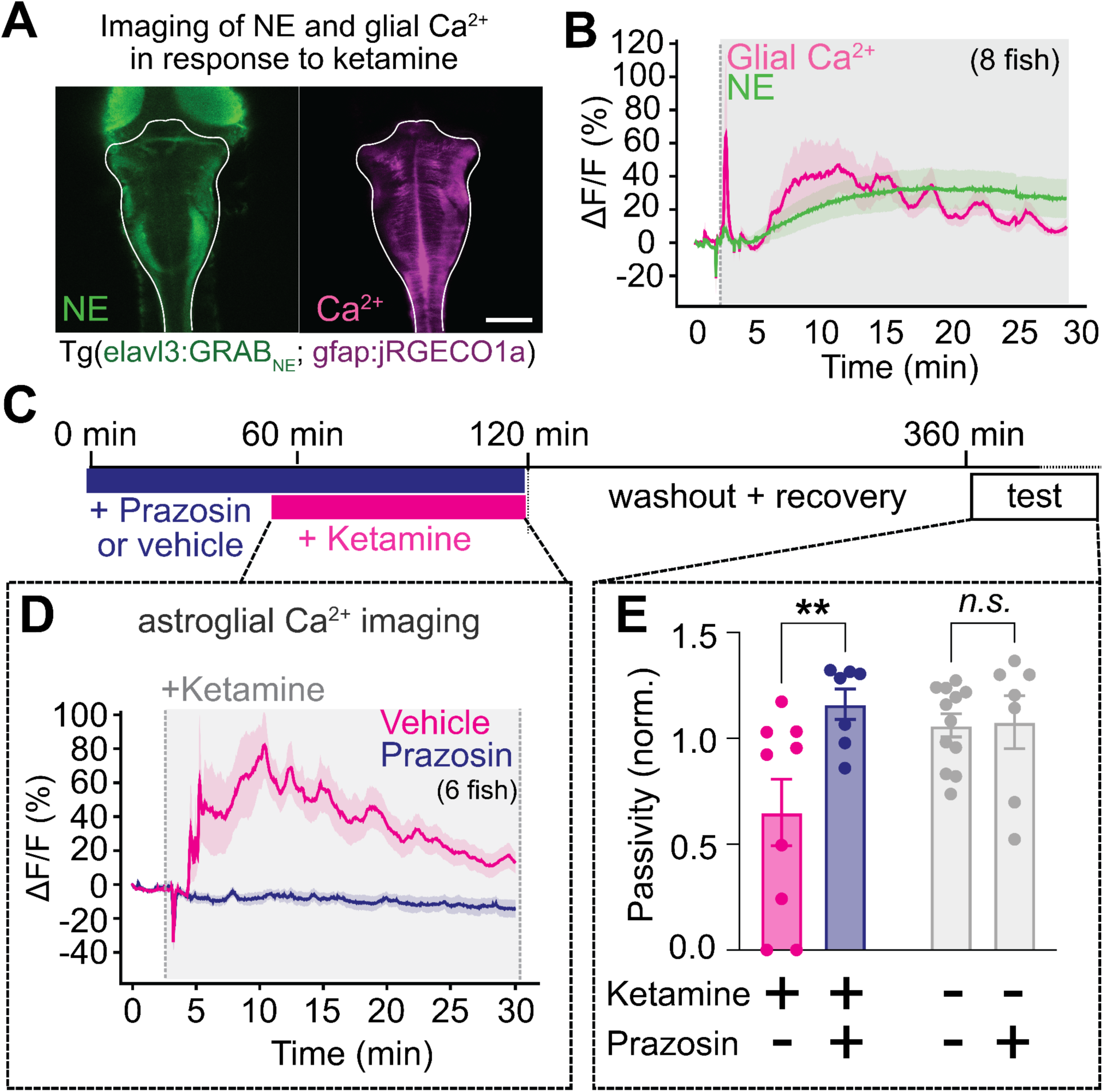
Noradrenergic signaling through α1-adrenergic receptors is required for ketamine’s action. (A) Fluorescence image of a larval zebrafish expressing GRAB_NE_ under the *elavl3* promoter (green) and jRGECO1a (glial Ca^2+^) under the *gfap* promoter (pink). Scale bar 100 μm. (B) Simultaneously imaged intracellular glial Ca^2+^ dynamics (jRGECO1a signal) and extracellular NE dynamics (GRAB_NE_ signal). Traces represent mean of fish (n = 8), shaded regions denote s.e.m. Shaded region indicates ketamine (200 μg/mL) in bath. Fluctuations at the onset of ketamine treatment are artifactual. (C) Schematic of experiments testing whether ketamine’s effects on astroglial Ca^2+^ and futility-induced passivity behavior depend on NE signaling. (D) Astroglial Ca^2+^ (jRGECO1b signal) response to ketamine (200 ug/mL) application in fish treated with 50 μM prazosin (n = 6 fish) or vehicle control (n = 5 fish). Control fish exhibit a stereotypical ketamine-driven astroglial Ca^2+^ elevation (orange), which is abolished in prazosin-treated fish (purple). (E) Open-loop passivity in our assay for fish treated with ketamine, with or without α1 adrenergic blocker prazosin (100 μM). Incubation with prazosin blocks the effect of ketamine in our assay. All error bars denote s.e.m. * p < 0.05, ** p < 0.01, *n.s.* p > 0.4. ΔF (z-scored) for each fish was calculated as the difference between mean total fluorescence before treatment subtracted to total fluorescence at a specific time point, divided by the standard deviation of total fluorescence before treatment, for hindbrain mask (D). Two-way ANOVA with Sidak’s multiple comparison test. N = 7 fish (control-prazosin and ketamine-prazosin), 9 fish (ketamine-vehicle), 11 fish (control-vehicle). p = Interaction (0.0140), 0.0057 (ket-prazosin vs ket-vehicle), 0.9947 (control-vehicle. vs control-prazosin).

### Prolonged opto/chemogenetic astroglial calcium elevation reduces futility induced passivity

To assess whether the NE-dependent astroglial calcium elevation induced by ketamine is sufficient to suppress passivity, we induced long-lasting calcium elevation in astroglia using two different methods. First, we optogenetically activated noradrenergic neurons using the *Tg(dbh:KalTA4; UAS:CoChR-eGFP)* fish line^38^, which expresses the channelrhodopsin CoChR (**Figure 5A**) under the *dbh* promoter. Continuous optogenetic activation of NE neurons with blue light (488 nm) induced a minutes-long calcium rise in hindbrain radial astroglia similar to the increase in calcium observed with ketamine, but with a much faster decay rate (**Figures S5B-S5D**; *Tg(dbh:KalTA4; UAS:CoChR-eGFP; gfap:jRGECO1a)* fish). Continuous stimulation of NE neurons for 30 minutes, followed by an hour of recovery, led to decreased futility-induced passivity (**Figure 5B**).

**Figure 5.**
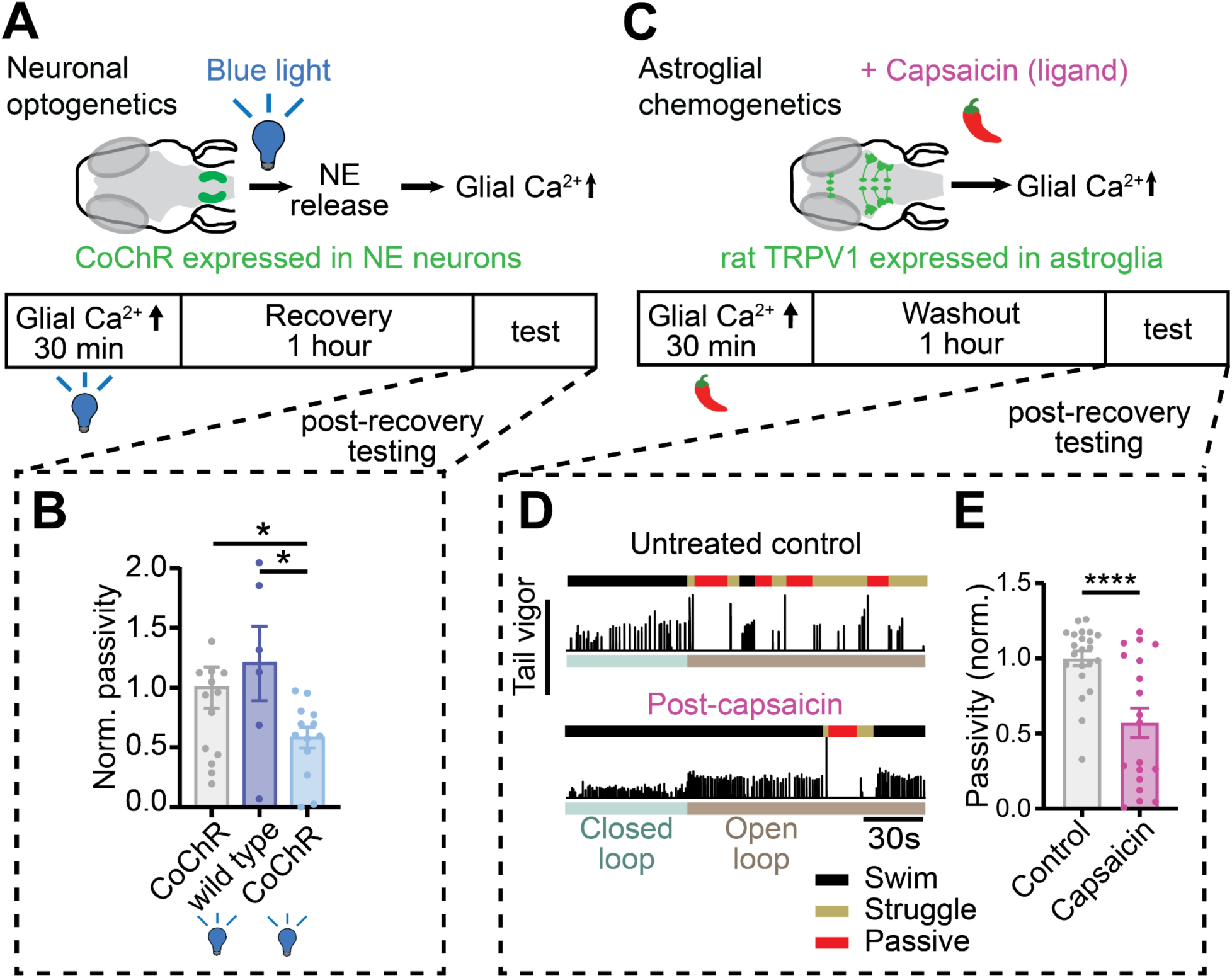
Elevating astroglial calcium recapitulates ketamine’s behavioral effects. (A) Timeline for experiments testing effects of stimulating glial calcium by activating norepinephrinergic (NE) neurons. *Tg(dbh:KalTA4);Tg(UAS:CoChR-eGFP)* fish expressing the channelrhodopsin CoChR in norepinephrinergic (NE) neurons were continually stimulated optogenetically with blue (488 nm) light for 30 minutes. Fish were allowed to recover for an hour then assayed for futile swimming. (B) Passivity normalized to clutch, non-stimulated controls. Optogenetic stimulation suppressed futilit-induced passivity following recovery compared to non-stimulated clutch controls and non-expression controls stimulated with blue light. Two-tailed t-test. N = 15 (dbh:CoChr, no light), 6 (wild type, blue light), 13 (dbh:CoChr, blue). p = 0.0197 (WT-Blue light vs DBH-Blue light). (C) Timeline for experiments testing effects of stimulating glial calcium directly through chemogenetics, Transgenic fish expressing rat TRPV1 in astroglia were treated with capsaicin for 30 min. After 30 min, capsaicin was washed out and fish allowed to recover for 1 h before being tested in our assay. (D) Example swimming of capsaicin-treated fish (bottom) and untreated clutch controls (top). Black, yellow, and red segments above swim vigor trace denote regular swimming, struggling, and passivity, respectively. (E) Passivity normalized to untreated clutch controls. Capsaicin-treated fish exhibit decreased futility-induced passivity, although the distribution of passivity duration is bimodal. Mann-Whitney test. N = 22 fish (Control), 19 fish (Capsaicin), p = 0.0005.

Because NE acts on neurons in addition to glia, we pursued more specific glial activation by chemogenetically stimulating astroglia in zebrafish that express the rat transient receptor potential V1 channel (TRPV1) under the *gfap* promoter (**Figures 5C** and **S5E**; Methods). Native TRPV1 in zebrafish is not sensitive to the chemical capsaicin, but rat TRPV1 is activated by its natural ligand capsaicin and induces an influx of calcium in radial astrocytes^38,70^. We found that exposure of fish expressing TRPV1 in astroglia to capsaicin for 30 minutes caused a long-lasting astroglial calcium elevation (**Figure S5F**) and suppressed futility-induced passivity after washout of the ligand (**Figures 5D** and **5E**). Pharmacological blockade of α1-AR had no effect on capsaicin-induced suppression of passivity (**Figure S5G** and **S5H**), indicating that the behavioral effects of astroglial calcium elevation occur downstream of α1-AR activation by norepinephrine. Thus, long-lasting calcium elevation in astroglia similar to that induced by ketamine is sufficient to suppress futility-induced passivity.

### Persistent suppression of neuronal and astroglial futility response following ketamine

While acute surges of astroglial calcium during futile swimming drive passivity^38^, the aftereffects of much larger and longer elevations evoked by the application of ketamine, and by our opto/chemogenetic manipulations, suppress passivity following recovery. To reconcile the seemingly contradictory effects of astroglial calcium during futile swimming and during ketamine administration, we hypothesized that although ketamine drives cytosolic calcium elevation when administered, when astroglial calcium later returns to baseline following washout and recovery, astroglia and potentially neuronal subtypes are less responsive to futile swims. To identify regions with long-lasting activity changes following ketamine exposure, we performed whole-brain light-sheet imaging of neuronal and astroglial calcium activity in *Tg(elavl3:H2B-jGCaMP7f; gfap:jRGECO1a)* fish after recovery from exposure to ketamine or vehicle for extended periods after drug washout (**Figure 6A**). We began by analyzing the responsiveness of hindbrain astroglia to open-loop swimming, because hindbrain astroglia are a critical node that integrates futility signals and drives passivity^38^. To control for differences in swim vigor, which might affect neural and astroglial responses, we analyzed swim-triggered responses in only high-vigor, struggle-like swims during open-loop (**Figures S6A and S6B**). Imaging revealed that, while the struggles analyzed pre-and post-ketamine treatment had similar vigors (**Figures S6C and S6D**), astroglial calcium responses to futile swimming were significantly diminished in ketamine-treated fish relative to controls (**Figures 6B-6D**). Because normally astroglial calcium surges trigger the passive behavioral state^38^, this decrease in the glial calcium response to futile swims provides a possible explanation for the decline in futility-induced passivity.

**Figure 6.**
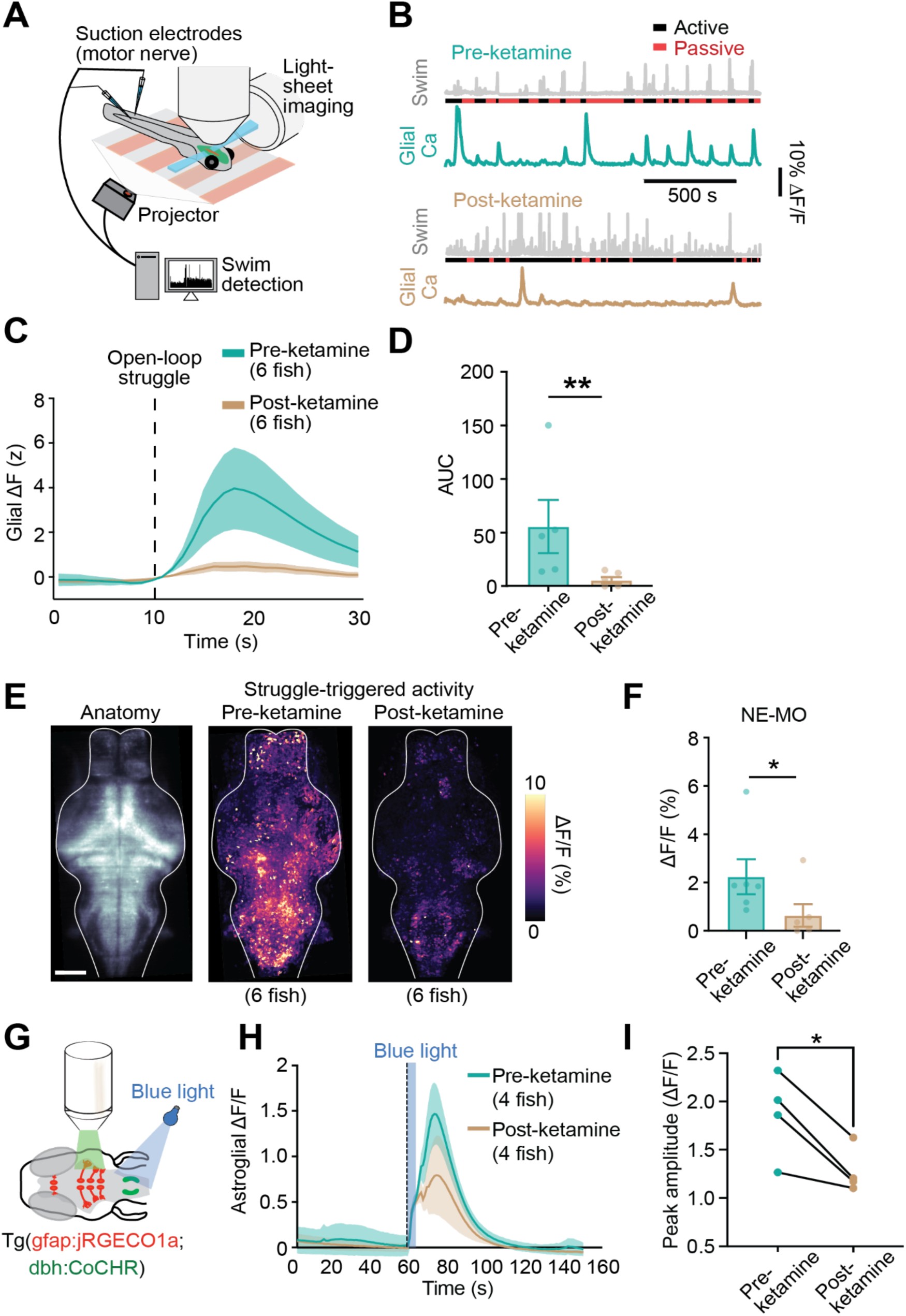
Persistent suppression of futility-activated neuron-astroglial circuits following ketamine exposure. (A) A light-sheet microscope recorded from most radial astrocytes and neurons in the brain at cellular resolution, using simultaneous dual-color nuclear H2B-GCaMP7f and jRGECO1a recordings, while fictive behavior was monitored and visual stimulus was delivered, as described in the Methods. (B) Swim traces (gray) showing active and passive periods and simultaneously recorded hindbrain astroglial calcium signals from two example *Tg(gfap:jRGECO1a)* fish either treated with vehicle control (pre-ketamine, top) or ketamine (post-ketamine, bottom). In both cases, fish were allowed to recover for 1 hour following washout before recordings were performed. (C) Average open-loop struggle-triggered jRGECO1a signal in hindbrain glia before and after recovery from ketamine treatment. (D) Area under the curve (AUC) for jRGECO1a signal following struggles in fish before and treatment with ketamine. Mann-Whitney test. N = 5 fish (pre), 6 fish (post). p = 0.0087. (E) (left) Anatomical max projection of *Tg(elavl3:H2B-GCaMP7f)* fish imaged during behavior. (middle) max projection of futile-swim triggered neural activity of untreated fish (N = 6 fish). (right) max projection of futile-swim triggered neural activity of ketamine-treated fish (N = 6 fish). (F) Average struggle-triggered ΔF/F in NE-MO cells before and after ketamine treatment. Mann-Whitney test. N = 6 fish (pre), 6 fish (post). p = 0.041. (G) Transgenic fish expressing the channelrhodopsin CoChR in norepinephrinergic (NE) neurons and the calcium reporter jRGECO1a in glia, were briefly stimulated optogenetically with blue (488 nm) light for 5s, while recording glial activity. (H) Average ΔF/F traces showing glial calcium responses after NE-MO stimulation with blue light, before (pre-ket) and after (post-ket) ketamine treatment, with different fish for pre/post. (I) Peak amplitude (ΔF/F) following blue light stimulation in the same pre-ket and post-ket fish. Two-tailed paired t-test. N = 4 fish pairs. p = 0.0268. All error bars denote s.e.m. * p< 0.05, ** p < 0.01.

A decrease in astroglial response to futility could arise from weaker noradrenergic futility signals from upstream neurons, or from an astroglia-autonomous decrease in sensitivity to the futility signals caused by, for example, noradrenergic receptor internalization. To investigate the contribution of decreased futility-triggered neural activity, we analyzed brain-wide neural activity signals corresponding to the swims and astroglial calcium events shown in **Figures S6A, S6B, 6B**, and **6C**. There was a significant decrease in futile struggle-triggered responses across the brain, including in NE-MO, a brainstem noradrenergic region that relays futility information to radial astroglia (**Figures 6E and 6F**). To determine if astroglia exhibited diminished sensitivity to NE after ketamine exposure, we assessed their response to selective activation of noradrenergic neurons using strong optogenetic stimulation which aimed to drive maximal firing of NE neurons (**Figure 6G**). We found that astroglial calcium responses to NE neuron stimulation were suppressed for at least one hour after ketamine washout (**Figures 6H and 6I**). However, this experiment is unable to rule out the possibility that NE release, even at maximal noradrenergic neuron firing, is diminished following ketamine treatment. To assess this possibility, one would need to image both NE and astroglial calcium while stimulating NE release, an experiment that is currently technically infeasible due to the incompatibility of our green NE sensor and blue-light channelrhodopsin. Therefore, we conducted an additional experiment using a different method (mild electric shock^71^) to trigger NE release from the locus coeruleus^71^ (**Figure S6E**). In control fish, mild electric shock induced an initial strong increase in astroglial calcium, while in ketamine treated fish, electric shock evoked a diminished increase in astroglial calcium (**Figure S6F**), Thus, just as with optogenetic stimulation of NE neurons, the astroglial response to mild electric shock was on average blunted by ketamine. To determine whether this decrease in astroglial calcium response is due to a decrease in NE release, we repeated the same experiments in fish expressing GRAB_NE1.0_. There was no difference in shock-evoked NE release between control and ketamine-treated animals (**Figure S6G**), suggesting that ketamine induces a long lasting decrease in astroglial sensitivity to NE.

### Ketamine induces a long-lasting increase in astrocyte calcium activity in mice

The stimulatory effect of NE on astroglial calcium is evolutionarily conserved; NE and homologous molecules directly increase astroglial calcium signaling in flies^21^, fish^38,71^, and mammals^22–24,26^. Because ketamine has been observed to affect NE levels in mammalian brains^16^, potentially related to what we observed in the larval zebrafish, ketamine may similarly alter the dynamics of astrocyte calcium signaling in the mammalian CNS. However, a previous study showed that the ketamine/xylazine combination used routinely for anesthesia in mice suppresses astrocytic calcium levels^72^. Because xylazine is known to suppress noradrenergic signaling through its activation of auto-inhibitory ɑ2 adrenergic receptors, we re-examined the effect of ketamine on mammalian astrocytes under non-anesthetic conditions.

We monitored intracellular calcium levels in astrocytes of *Aldh1l1-CreER;R26-lsl-GCAMP6s* mice using two photon microscopy in awake, unanesthetized conditions (**Figure 7A**, **Figure S7A**). Astrocytes were visualized through a chronic cranial window implanted above the retrosplenial cortex (RSC), a region implicated in producing the dissociative effects of ketamine^73^. Acute intraperitoneal injection of ketamine induced a striking, widespread increase in cytosolic calcium within astrocytes throughout the imaging field (**Figure 7B**; **Video S4**) not seen in mice injected with vehicle alone (**Figures S7B-S7E**). The response to ketamine peaked several minutes after injection and lasted for >10 minutes in all animals tested (**Figure 7C and 7D**; **Figures S7F-S7I**), similar to the time course seen in zebrafish. The astrocyte calcium elevation consisted of both a plateau response and an increase in discrete calcium transients within the field of view that waned with increasing time after injection (**Figure S7F-S7I**). These responses were abolished by treatment with dexmedetomidine, a highly specific α2 adrenoreceptor agonist that inhibits NE release (**Figure 7E**). Together, these data indicate that ketamine elicits a prolonged increase in astrocytic calcium by enhancing the activity of noradrenergic neurons. The presence of this phenomenon in both fish and mammals suggests that ketamine engages an evolutionarily conserved mode of astrocyte neuromodulation, consistent with the high conservation of this modulatory pathway from flies to humans^21–24,38^.

**Figure 7.**
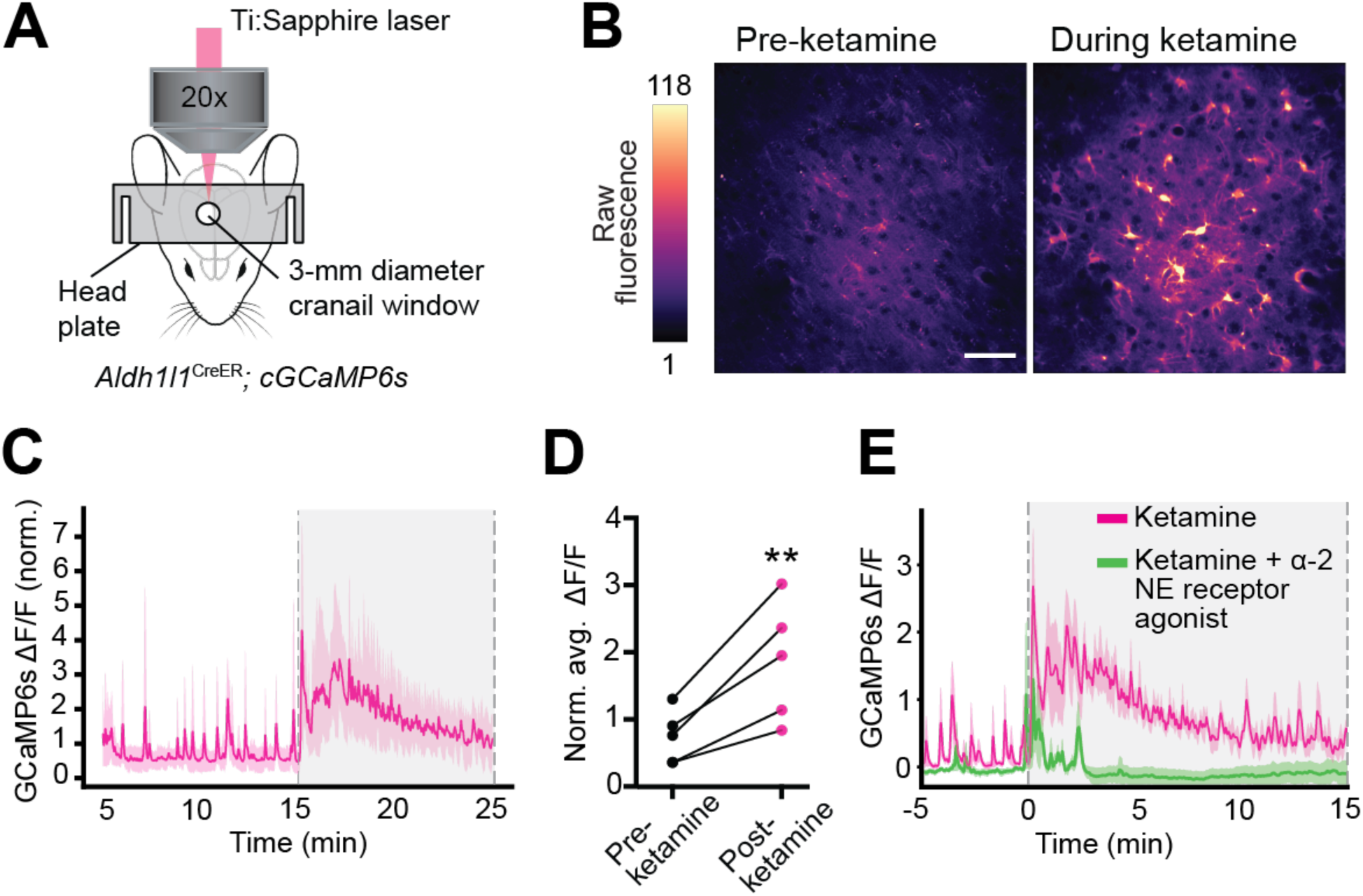
Ketamine elevates astrocytic calcium in mammalian astrocytes *in vivo* through conserved signaling pathways. (A) Schematic showing setup for *in vivo* imaging of cortical astrocytic calcium in awake mice. (B) Fluorescence micrographs of GCaMP6s signal in the retrosplenial cortex (RSC), for an example mouse at two time points illustrating elevation of cytosolic calcium following IP injection of 20 mg/kg ketamine. Scale bar 50 μm. (C) Average RSC fluorescence trace for mice injected with 20 mg/kg ketamine (N=5 mice) at 15 min. (D) Quantification of RSC fluorescence change before and after ketamine injection, normalized to control. Paired two-tailed test. N = 5 mice. p = 0.0089. (E) Average RSC fluorescence trace for mice injected with 20 mg/kg ketamine or 20mg/kg ketamine plus an alpha-2 agonist, dexmedotomidine. N = 5 mice. ** p < 0.01.

## Discussion

An expansive body of research has documented the myriad effects of ketamine on neuronal physiology potentially critical for its antidepressant effects; however, this complexity has confounded attempts to develop a cellular and circuit-based understanding of how brief ketamine exposure leads to long lasting antidepressant effects. Our studies provide evidence that a significant driver of this phenomenon occurs through neuromodulatory regulation of calcium levels in astroglia. Emerging evidence has suggested that astroglia play important and evolutionarily conserved roles in brain state modulation^20,21,24,26,71^. Our work links astroglial modulation of brain state with previous work on the role of astroglia in both depression and the efficacy of antidepressants. Ketamine has been shown to drive monoaminergic signaling to elevate NE and 5-HT in the extracellular space^16,62^, and, in the case of serotonin^74^, this elevation has been shown to be important to mediate antidepressant-like effects of ketamine in rodents. Astrocytes have been shown to respond to elevations in NE^21–24,38^ and serotonin^31^ with elevations in cytosolic calcium. Furthermore, astroglial responsiveness to serotonin has recently been shown to be disrupted in rodent depression models, and stimulating prefrontal cortex astrocytes chemogenetically has been shown to rescue depression-related behavioral phenotypes^31^. Ketamine has also been shown *in vitro* to decrease astroglial calcium responses to stimuli like ATP and to attenuate calcium-coupled secretion of chemical messengers by astroglia^75^. Our results suggest that profound astroglial calcium elevation in response to the ketamine-evoked enhancement of monoaminergic signaling is key to its ability to suppress futility-induced passivity. One possible mechanism by which the profound and long-lasting NE elevation caused by ketamine could lead to a decreased astroglial response to NE following washout is receptor desensitization through phosphorylation^76^, but future studies should explore additional potential mechanisms.

A fundamental challenge is faced by efforts to develop pharmacological interventions for psychiatric disorders and understand the mechanisms underlying their efficacy. Namely, while the proximal targets of pharmacological interventions are proteins (e.g. receptors or channels on neurons or other cell types), the ultimate determinant of the intervention’s efficacy is whether it can correct behavioral abnormalities. The mapping from pharmacological perturbation and behavioral change is highly non-linear, as it involves changes in cellular activity across many brain regions. Therefore, the field requires models in which the activity of neural and non-neuronal cells, and other relevant variables like neuromodulator levels, can be measured before, during, and after pharmacological intervention and simultaneously with behavior. Our work illustrates, using the fast-acting antidepressant ketamine as an example, the utility of the larval zebrafish as one such model. The larval zebrafish has long been used for drug discovery and behavioral profiling in high-throughput assays for simple behavioral phenotypes^35^. Our study demonstrates that larval zebrafish can also be used to assay complex behavioral phenotypes, such as futility-induced passivity. Furthermore, the optical and genetic accessibility of the larval zebrafish presents a unique opportunity to investigate the effects of antidepressants and other pharmacological interventions on both neurons and non-neuronal cells in the brain of a behaving animal. Because larval zebrafish survive and behave for many hours following head-fixation, the effects of these compounds can be studied over many timescales. In addition, because the entire body of the fish is transparent, it could serve as a model for the effects of compounds on the peripheral nervous system and other organ systems, in the future, to identify the physiological mechanisms by which the many side effects of pharmacological agents, including antidepressants, are made manifest^77,78^.

Many of the circuits and cell types that modulate affective state, motivation, and stress responses are evolutionarily ancient, and the molecular targets of many pharmacological agents, including ketamine and other antidepressants, are similarly conserved. Therefore, observations made in the larval zebrafish could generalize to mammalian systems and inform further study in mammals. Indeed, we found that ketamine causes profound and long-lasting calcium elevations in the mouse cortex through an evolutionarily ancient NE-to-astrocyte signaling axis. Astrocytes throughout the mammalian brain, including in the hindbrain^24^, express α1-ARs and respond to NE, suggesting that ketamine may exert widespread increases in astrocyte calcium in the brain. Astrocytic calcium is also elevated by transcranial direct current stimulation (tDCS) in rodents^79^, and the antidepressant effects of tDCS require many of the same molecular components (NE and astrocyte α1-ARs) as the astroglial futility pathway in larval zebrafish^38^, suggesting that diverse antidepressant interventions may act in part through shared action on evolutionarily conserved neuromodulatory systems spanning neurons and astrocytes.

While we investigated ketamine’s effects on cellular activity 1-2 h following washout, ketamine exerts potent behavioral effects for days to weeks in rodents and humans^1,11^. Some of ketamine’s very long-lasting effects could be explained by changes in neuronal plasticity^11,15,17,80^. Proteins secreted by astroglia, such as hevin^81^, are critical mediators of synaptogenesis. Interestingly, recent work has implicated both astrocyte calcium and hevin in cocaine-induced plasticity^82^. Ketamine might act through similar pathways to elicit the neuroplasticity important for its antidepressant effects. Understanding how the same astroglia-mediated synaptogenetic pathways can underlie the pathophysiology of some agents and the therapeutic properties of others remains an important open question.

In addition to synaptic plasticity, the profound calcium elevation in astrocytes may cause long-lasting changes in astroglial physiological properties that contribute to their days-long modulation of behavior. For example, changes in astroglial Kir4.1 potassium channel expression have been shown to cause lateral habenula bursting contributing to behavioral changes in rodent models of depression^69^. While ketamine acutely inhibits lateral habenula bursting through its blockade of NMDARs^13^, and this blockade may persist for up to 24 hours due to channel trapping of the drug^68^, bursting inhibition lasting for much longer than an hour may occur through calcium-mediated changes in membrane insertion of Kir4.1 channels, as has been shown *in vitro*^83^. Our studies suggest that the potent effects of ketamine may reflect a previously unrecognized synergy between NE modulation of astrocyte calcium and direct NMDAR inhibition^13^ on neurons and potential parallel pathways for mediating futility-induced state switching.

Our results connect the fields of glial biology and the neurobiology of depression. Astroglial responsiveness to futility is blunted after cytosolic calcium returns to baseline following ketamine washout, suggesting that this drug induces prolonged changes in astroglial physiology, providing a mechanism through which the physiological effects of ketamine impacts circuit function and behavior. Astrocytes express receptors for many classes of signaling molecules^84^, but the functional and behavioral consequences of astroglial receptor activation have remained unclear. Our studies here show that astroglial activation can have profound, lasting effects on behavioral and neural circuit state. Understanding how fast-acting antidepressants act on non-neuronal cell types, in addition to neurons, could lead to new insight into the cellular and circuit pathways that go awry during major depressive disorder to inform the discovery and rational design of new fast-acting, effective therapeutics.

## Acknowledgements

We would like to thank Dr. Bryan Roth and Dr. Jon Ellman, and their labs, for providing us with R-69. We would like to thank Herriet Hsieh for discussions and comments on the manuscript. We would also like to thank Sabrina Boutselis for designing the red hot chili pepper illustration. We would like to thank all laboratory members for feedback, Mark Ellisman for discussions, and Brett Mensh for advice on the manuscript.

## Funding

Boehringer Ingelheim Fonds Graduate Fellowship (MD, AR)

German Research Foundation SPP1757 SA2114/2 (GS)

Howard Hughes Medical Institute (ABC, MD, SN, MBA)

NIH Grant 1R01NS126043 (AEC, SB)

NIH Grant P30 NS050274 (DEB)

NIH Grant P50 MH084020 (DEB)

NIH Grant 1R01GM128997 (DEO)

NIH Grant R35 NS122172

NIH Grant U19NS104653 (FE)

NIH Grant 1R01NS124017 (FE)

NSF Grant IIS-1912293 (FE)

NSF GRFP DGE1745303 (ABC)

NSF GRFP DGE2139757 (EH)

Simons Foundation SCGB 542943SPI (FE, MBA)

## Author contributions

Conceptualization: MD, ABC, MCF, MBA

Methodology: MD, ABC, EH, SN, SB, GS, AR, DAP

Investigation: MD, ABC, EH

Visualization: MD, ABC, EH

Funding acquisition: AEC, DEO, DAP, DEB, MCF, FE, MBA

Project administration: MCF, FE, MBA

Supervision: AEC, DEO, DEB, MCF, FE, MBA

Writing – original draft: MD, ABC

Writing – edit and revision: MD, ABC, EH, AEC, DEO, DEB, MCF, FE, MBA

## Declaration of interests

DEO is a co-founder of Delix Therapeutics, Inc., serves as the Chief Innovation Officer and Head of the Scientific Advisory Board, and has sponsored research agreements with Delix Therapeutics. Delix Therapeutics has licensed technology from the University of California, Davis. AEC is a co-founder of Q-State Biosciences. All other authors declare that they have no competing interests.

## Data and materials availability

All plots were generated using Prism (GraphPad); data used to generate all figures as well as Prism files used to generate them will be deposited on GitHub (https://github.com/alexbchen). Jupyter notebooks (Python 3.7) were used to process raw data. Raw data will be made available by the corresponding author upon request. Python and C++ code used will be made available by the corresponding author upon request. Fish lines will be made available upon request.

## Materials and Methods

### Experimental model and subject details

Experiments were conducted according to the guidelines of the National Institutes of Health and were approved by the Standing Committee on the Use of Animals in Research of Harvard University. Animals were handled according IACUC protocols #1836 (Prober lab), #2729 (Engert lab), 18-11-340-1 (Fishman lab), and 22-0216 (Ahrens lab). For all experiments in larval zebrafish, we used wild-type larval zebrafish (strains AB, WIK or TL), aged 5–8 days post-fertilization (dpf). We did not determine the sex of the fish we used since it is indeterminate at this age. Fish were raised in shallow Petri dishes and fed ad libitum with paramecia after 4 dpf. Fish were raised on a 14 h:10 h light:dark cycle at around 27°C. All experiments were done during daylight hours (4–14 h after lights on). All protocols and procedures were approved by the Harvard University/Faculty of Arts and Sciences Standing Committee on the Use of Animals in Research and Teaching (Institutional Animal Care and Use Committee), and the Janelia Institutional Animal Care and Use Committee.

For mouse experiments, female and male adult mice were used and randomly assigned to experimental groups. All mice were healthy, and none were excluded from the analysis. Mice were maintained on a 12-hour light/dark cycle, housed in groups no larger than five, and received food and water ad libitum. All mouse experiments were conducted in accordance with the National Institute of Health Guide for the Care and Use of Laboratory Animals and protocols approved by the Animal Care and Use Committee at Johns Hopkins University.

### Fish lines

For all behavioral experiments we used:

*Wild type - strains AB and WIK*

For imaging of astroglial calcium we used:

Cytosolic, red calcium indicator. *Tg(gfap:jRGECO1a)*^38,85^

For imaging of neuronal calcium we used:

Nuclear-localized, green calcium indicator. *Tg(elavl3:H2B-jGCaMP7f)^jf^*^90^ ^86^

For optogenetic activation of norepinephrinergic neurons we used:

Channelrhodopsin expressed under *dbh* promoter. *Tg(dbh:KalTA4);Tg(UAS:CoChR-eGFP)*^87^

For chemogenetic activation of astroglia we used:

Rat transient receptor potential cation channel subfamily V member 1 (TRPV1) expressed under *gfap* promoter. *Tg(gfap:TRPV1-T2A-eGFP)^jf^*^64^ (*this paper*)

For imaging extracellular norepinephrine concentrations we used:

GRAB_NE1.0_^65^ green fluorescent extracellular norepinephrine sensor expressed under *elavl3* promoter. *Tg(elavl3:GRAB_NE_2h)* (*this paper*).

For imaging extracellular serotonin concentrations we used:

GRAB_5-HT4.0_ green fluorescent extracellular serotonin sensor expressed under *elavl3* promoter. *Tg(elavl3:GRAB_5-HT4.0_)* (ZFIN ct876Tg) (*this paper*).

### Zebrafish transgenesis

We generated the *Tg(gfap:TRPV1-T2A-eGFP)^jf^*^64^*, Tg(elavl3:GRAB_NE1.0_2h)*and *Tg(elavl3*:*GRAB_5-HT4.0_)* lines used in this paper. The lines were generated in casper background^88^ using the Tol2 method^89^.

### Embedding of larval zebrafish for tail-tracking experiments

Larval zebrafish aged 6-8 dpf were embedded in small round Petri dishes (e.g. Corning #351006); importantly, dishes should not be tissue culture-treated to allow agarose to adhere. Solutions of 2% low melting-point agarose (Sigma-Aldrich A9414) were prepared by heating powdered agarose in system water and agitating until solution was clear. The 2% agarose solution was kept at 42-48 degrees Celsius. To embed fish, a small amount of 2% agarose solution was pipetted in the middle of a Petri dish. A larval zebrafish was then transferred, using either a small glass or Pasteur pipette, taking care to minimize addition of water to the agarose solution during transfer. Using either small forceps or a small (∼10-100 µL) plastic micropipette tip, larval zebrafish were gently rotated until they were dorsal side up. Fish often struggle when transferred to the agarose solution, so care should be taken to keep fish righted until agarose solidifies. Once the agarose solidified, we freed the tail of the fish by carefully cutting away the hardened agarose around the tail with a micro-scalpel (Fine Science Tools 10315-12). Care should be taken to remove enough agarose to prevent the tail from hitting or becoming stuck on agarose during swimming or struggle.

### Pharmacological treatment of larval zebrafish

Approximately 10-20 larval zebrafish aged 6-8 dpf were transferred to single wells in 12-well plates containing 2 mL of fish water (8-12 fish/well). Quantities of either the experimental pharmacological compound (Table S1) or vehicle control were added and fish were incubated for one hour. Fish were then washed three times in a larger Petri dish and then allowed to recover for one hour, following which experimentation would begin. For the MS-222 experiments, to avoid pH effects, the solution was buffered with 10mM HEPES and pH adjusted to 6.8-7.4 using a 1M solution of NaOH.

### Tail-tracking of embedded larval zebrafish and visual stimulus

For all behavioral experiments involving embedded larval zebrafish, we used a previously published, custom-build behavioral rig and custom-written code^91^. Briefly, we illuminated the fish and its environment using infrared light-emitting diode panels (wavelength 940 nm, Cop Security). The tail-posture of the fish was tracked using a camera (Grasshopper3-NIR, FLIR Systems) with a zoom lens (Zoom 7000, 18–108 mm, Navitar) and a long-pass filter (R72, Hoya). We determined the posture of the tail, as well as its tip angle, by analyzing the position of ∼25 equally spaced, user-defined key points along its length. Posture was determined and recorded in real-time at 90 Hz using custom-written Python scripts (Python 3.7, OpenCV 4.1).

### Tracking of freely swimming larval zebrafish

For all behavioral experiments involving freely-swimming larval zebrafish, we also used a previously published, custom-build behavioral rig and custom-written code ^91^. The illumination and detection of freely swimming fish used the same behavioral rigs as described for embedded fish. To track the position of fish and determine swim bouts in real time at 90 Hz, we used custom-written Python scripts (Python 3.7, OpenCV 4.1). The background of the camera image was subtracted and the body of the fish identified by center of mass. Orientation was determined as the axis of largest pixel variance in the identified body. Swim bouts were detected by computing a 50-ms rolling variance and identifying thresholded peaks.

### Passivity computation in zebrafish

Passive periods were operationally defined to be periods greater than 10 s in length in which the fish did not perform a swim bout. To determine the percentage of the open-loop interval spent passive per trial, the total length of all passive periods within the open-loop interval of a trial was summed and the sum divided by the length of the open-loop interval. The same computation was performed to determine passive fraction for closed-loop intervals as well.

### Epifluorescence imaging of neurons and radial astrocytes in embedded larval zebrafish

Larval zebrafish aged 6-8 dpf were embedded in small round petri dishes and their tails freed as described previously. To image neurons, we used previously published fish lines (see Fish Lines section) *Tg(elavl3:H2B-GCaMP7f)* for neural imaging and *Tg(gfap:jRGECO1a)* for astroglial imaging. Imaging was performed using a dissecting microscope (Olympus MVX10) with a CMOS camera (IDS Imaging UI-3370CP-NIR) and an LED lamp for fluorescent imaging (X-Cite 120 LED mini). For analysis, each video frame was registered to a reference brain using OpenCV and fluorescence in each defined brain region was extracted with manually segmented masks. Fluorescence Z-score was calculated as the difference between mean total fluorescence before treatment subtracted to average baseline fluorescence, divided by the standard deviation of total fluorescence before treatment, for each mask.

### Confocal imaging of extracellular norepinephrine and astroglial calcium in zebrafish

Larval zebrafish aged 6-8 dpf of the line *Tg(elavl3:GRAB_NE_2h);Tg(gfap:jRGECO1a)* were embedded in small round petri dishes and their tails freed as described previously. Two-color imaging was performed at 0.5 Hz using a Zeiss LSM 980 NLO confocal microscope (488 and 561 nm laser illumination) with a 20x magnification objective (Zeiss #421452-9800-000). Ketamine (200 mg/L) was added to the bath with a 200 uL micropipette at 100 frames. For experiments in which prazosin was added prior to imaging, fish were incubated in prazosin (100 μM in bath) for at least 2 h before embedding. For experiments in which MS-222 was added prior to imaging, fish were incubated in MS-222 (150 mg/L) for at least 30 min prior to imaging. Fluorescence was extracted with manually segmented masks. For each fish, the same mask was used for the green and red channels. DFF calculations were performed as described above.

### Fictive behavior in virtual reality and light-sheet imaging in zebrafish

The setup used for fictive behavior during light-sheet imaging was adapted from Mu et al. 2019. Animals in fictive-behavior experiments for light-sheet imaging were not paralyzed. Fish were directly transferred to an acrylic pedestal in a custom-fabricated sample holder with glass windows allowing side illumination and immobilized with 2% agarose. Agarose was removed by hand over a small square in the tail to allow access for the electrodes while still restricting overall movement, and agarose was removed around the head to minimize aberration of the light-sheet illumination of the brain. One suction electrode (∼60 μm inner diameter) filled with external solution, was placed over the dorsal side of the fish’s tail and attached with gentle negative pressure. The voltage signal recorded by this tail was amplified and filtered (band-pass 300 Hz - 3 kHz) with a MultiClamp 700B amplifier. This signal was then smoothed through convolution with an exponential filter and used as the ‘swim signal’.

For the struggle analysis performed in Figures 6 and S6, we first detected swims by detecting peaks in the swim signal (minimum inter-peak interval > 500 ms). We then z-scored swim amplitudes and manually set a z-score threshold for each fish, above which a swim would be classified as a struggle. The same set of z-score thresholds were used to analyze neural and glial activity.

### Analysis of zebrafish light-sheet calcium imaging data

We extracted cell segments from raw fluorescence data with a volumetric segmentation pipeline available from https://github.com/mikarubi/voluseg, and described previously^38,48^. We calculated ΔF/F for each cell segment using a 100s rolling baseline window as described in Yang et al., 2022^48^. For the acute ketamine imaging, we calculated the log2 fold change for each cell segment as:

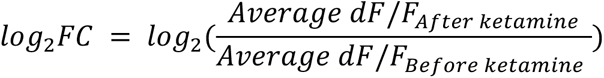

For the activation-inhibition maps, we sorted cell segments according to the fold change and plotted the top 10% most activated and inhibited segments, for each fish.

For the fictive behavior calcium imaging data, we ran the segmentation and obtained ΔF/F for each cell segment as described above.

### Registration of different zebrafish brain volumes and Z-Brain masks

To compare results across fish we registered all experiments to a representative brain volume, using BigStream (https://github.com/JaneliaSciComp/bigstream).

For the light-sheet calcium imaging data, after registration of all experimental volumes together, we registered a Z-Brain reference H2B-mCherry volume^59^ to the representative brain volume and applied the same transformation to previously defined reference masks (total of 294 masks) for every brain region.

### Optogenetic and chemogenetic activation experiments in zebrafish

For the optogenetic experiments, *Tg(dbh:CoChR-eGFP)*^87,92^ fish were incrossed and their larvae raised until 6-8 dpf. Fish were screened using an epifluorescence scope and selected according to the GFP signal in NE-MO cells, and raised to 6-8 dpf. Positive fish were then placed on top of a 16x16 blue LED matrix and stimulated for 30 minutes with blue light (0.3W, Newegg #9SIA9DCJ4V0029) or control ambient light. After 1h recovery, fish were assayed in our futility-induced passivity test. For the imaging experiments, positive fish also expressing *Tg(gfap:jRGECO1a)* were imaged in a confocal microscope and stimulated from the top using a blue light (450-460nm) LED source (UltraFire #WF-508BL) for 5 seconds.

For the chemogenetic experiments, *Tg(gfap:TRPV1-T2A-eGFP)^jf^*^64^ were crossed with WT fish from a casper background. Fish were screened using an epifluorescence scope and selected according to the GFP signal in hindbrain and spinal cord glia, and raised to 6-8 dpf. Positive fish were then incubated with 2µM capsaicin (Sigma #12084) in fish water (0.025% DMSO final concentration) or an equivalent concentration of vehicle for 20-30 minutes. Fish were then washed and placed in fresh fish water. After 1h recovery, fish were assayed in our futility-induced passivity test.

### Mild electric shock experiments in zebrafish

We glued a pair of electrodes to a petri dish using JB-Weld #8265S. The electrodes were separated by ∼3cm. The fish were placed perfectly in between the electrodes and parallel to the metal, embedded in agarose and tail-free. For stimulation, we used a 20V 200ms stimulus (SD9K stimulator, Grass Instruments) and performed simultaneous imaging of glial calcium activity in the epifluorescence microscope as described above.

### Tamoxifen preparation and induction in mouse

Conditional knockin Rosa26-lsl-GCaMP6s^93^ and Aldh1l1-CreER^94^ mouse lines have been previously described. To induce GCaMP6s expression in *Aldh1l1-CreER;GCaMP6s* transgenic mice, TAM (Sigma-Aldrich, T5648) was freshly prepared on the first day of the injection at 10 mg/mL in sunflower seed oil (Sigma-Aldrich, S5007) through intermittent sonication at room temperature (RT). Adult (> 6 weeks) mice were injected intraperitoneally (i.p.) with a dosage of 100 mg/kg body weight (b.w.) for five consecutive days, once per day. Every injection was at least 20 hours (hrs) apart. The remaining tamoxifen solution was stored at 4°C in the dark for a maximum of 5 days. All experiments were performed at least 2 weeks after the last tamoxifen injection.

### Head plate installation and cranial window surgery in mouse

The day before cranial window surgery, dexamethasone (VetOne, NDC#13985-037-02) was given through drinking water to the mice (1 mg/kg) that completed TAM administration (see above). The next day, animals were anesthetized with inhaled isoflurane (0.25-5%) and placed in a custom-made stereotaxic frame. Surgery was performed under standard and sterile conditions. After hair removal and lidocaine application (1%, VetOne, NDC 13985-222-04), the mouse’s skull surrounding the left retrosplenial cortex was exposed and the connective tissue was carefully removed. VetbondTM (3M) was used to close the incision site. A custom-made metal head plate was fixed to the cleaned skull using dental cement (C&B Metabond, Parkell Inc.). A 3-mm diameter circular craniotomy was then performed using a high-speed dental drill bit. The center of the craniotomy was located 1 mm lateral to lambda. Dura was left intact and the cranial window was then sealed with a custom-made 3 mm diameter circular #1 (0.17 mm) coverslip using VetbondTM. A layer of cyanoacrylate (Krazy Glue) was applied on top of the Vetbond to secure the coverslip. Animals recovered in their home cages for at least 2 weeks before imaging.

### In vivo 2P laser scanning microscopy and ketamine delivery for astrocyte Ca^2+^ activity in <u>mouse</u>

Two-photon laser scanning microscopy was performed with a Zeiss LSM 710 microscope equipped with a GaAsP detector, which uses a mode-locked Ti-Sapphire laser (Coherent Chameleon Ultra II) tuned to 920 nm. The head of the mouse was immobilized by attaching the head plate to a custom-made stage mounted on a vibration isolation table, and the body of the mouse was housed in a custom-made plastic restrainer. Images were collected 100 – 150 mm below dura using a coverslip-corrected Zeiss 20x/1.0 W Plan-Apochromat objective with a pixel dwelling time of 1.58 ms and scanning speed of 2 Hz. Mice were kept on the stage for no more than two hours and all *in vivo* imaging experiments were performed during the day.

For *in vivo* ketamine delivery, an I.P. catheter was inserted into the lower right quadrant of the animal’s abdomen (DDP Medical, Cat. 405311). Animals were then imaged as described above. Within 15-30 minutes of the initiating imaging, animals were injected intraperitoneally with ketamine (20 mg/kg; MWIAH; dissolved in 0.9% NaCl) or vehicle (0.9% NaCl). In experiments with α2-adrenergic agonist, 20 mg/kg ketamine was co-administered with dexmedetomidine (0.01 mg/kg; PRECEDEX).

### Quantification and statistical analysis of mouse imaging data

Statistical analyses were performed in GraphPad Prism. All statistical tests in this study were two-tailed and the details for each analysis are shown in each figure legend.

For the statistical analysis across atlas brain regions for Figure 2, we bootstrapped a 95% confidence interval in shuffled data by shuffling the brain area ID corresponding to each segmented unit, then taking the top 10% of shuffled units to calculate a mean activation for each brain region. This was repeated separately for all fish, then a mean taken across fish to complete a single bootstrap trial. We repeated the bootstrapping process 10,000 times, then took the 2.5 and 97.5 percentiles to generate the confidence interval.

## Supplementary Materials

Supplementary Figures S1-S7 Table S1

Supplementary Videos S1-S4

### Supplementary Figures

**Figure S1.**
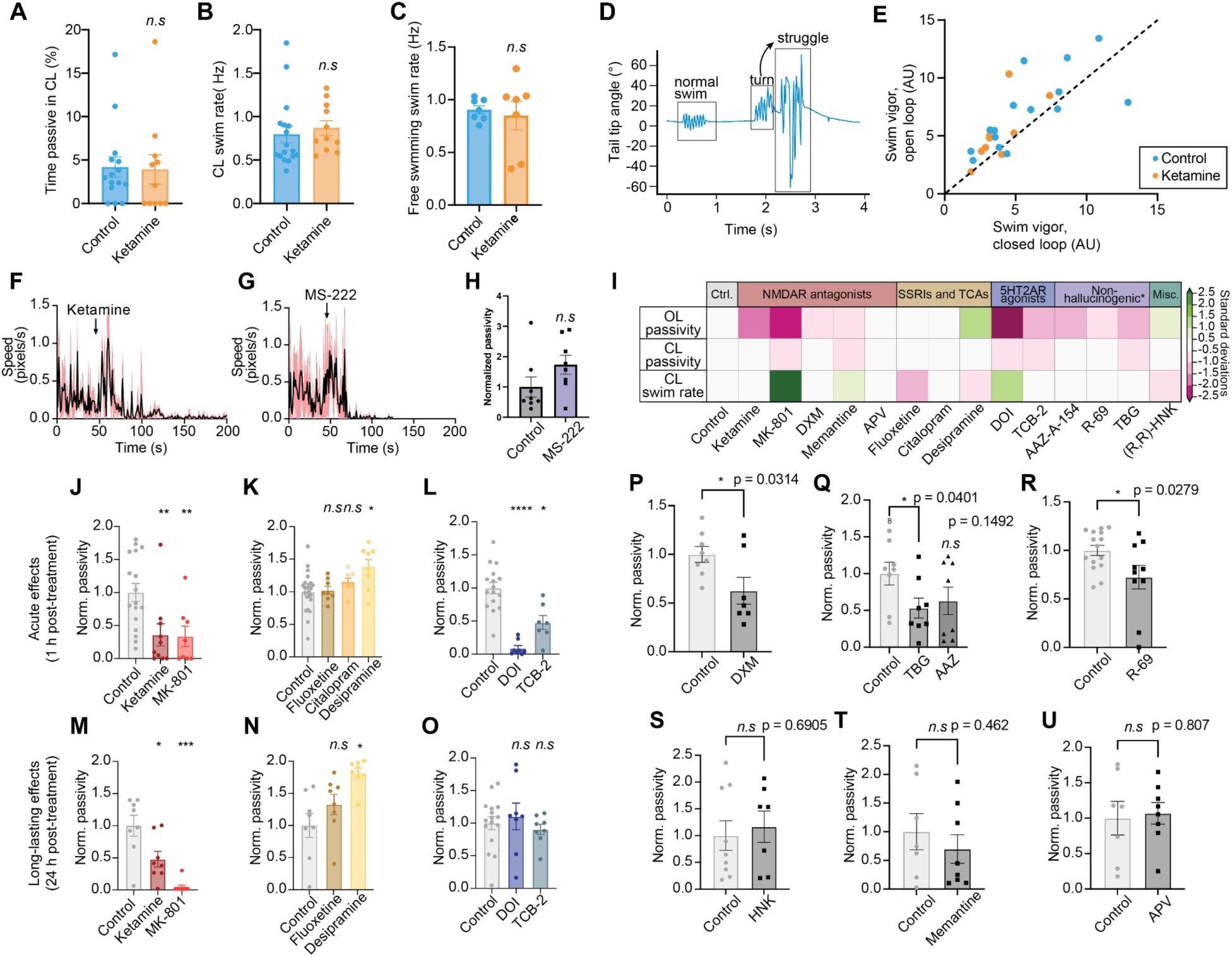
Relates to Fig. 1. (A) Average percentage of CL interval spent in the passive state. Individual dots represent trial averages for each fish. Control and ketamine fish exhibit similar average % passive time in CL (t-test, p = 0.8921). (B), A comparison of closed-loop (CL) swim rate in control and ketamine-treated fish. Individual dots represent trial averages for each fish. Control and ketamine fish exhibit similar average swim rate (t-test, p = 0.5898). (C), A comparison of closed-loop (CL) swim rate in control and ketamine-treated fish. Individual dots represent trial averages for each fish. Control and ketamine fish exhibit similar average swim rate (t-test, p = 0.70). (D) Example snippet showing the angle made by the tip of the tail relative to the fish’s long body axis during a 4 s period during open-loop swimming. Three different bout types, a representative normal swim, turn (asymmetric tail angle) and struggle (large amplitude, uncoordinated swimming), can be seen and are manually annotated. The observed turn transitions into a struggle with little inter-swim interval, denoted by the arrow. (E) Average swim vigor (peak value of detected swim bouts yielded by 500 ms moving standard deviation (see Methods) during open loop plotted against average swim vigor during closed loop. Each point represents the trial average of an individual fish. While the OL to CL ratio does not differ between control and ketamine-treated fish (t-test on slope, p = 0.7097), both control and ketamine-treated fish exhibited more vigorous swimming in OL than in CL (paired t-test mean vigor OL-CL>0, p = 0.0268 for ketamine, p = 0.0347 for control).(I), Open-loop passivity, closed-loop passivity, and closed-loop swim rate of fish treated with vehicle (control) or various different pharmacological compounds (normalized to mean of control) in the futility-induced passivity assay. Results are grouped by major pharmacological target of treatment compound. (J-L) Acute (1 h post-washout) effects of (J), Phencyclidine-site NMDA receptor antagonists, (K), 5-HT2A receptor agonists, and (L), selective serotonin reuptake inhibitors (SSRIs) and tricyclic antidepressants (TCAs) on open-loop passivity. Phencyclidine-site NMDA receptor antagonists and 5-HT2A receptor agonists exhibit acute passivity-suppressing effects, but SSRIs and TCAs do not. (M-O) Persistent (24 h post-washout) effects of (M), Phencyclidine-site NMDA receptor antagonists, (N), 5-HT2A receptor agonists, and (O), SSRIs and TCAs on open-loop passivity. Only the phencyclidine-site NMDA receptor antagonists exhibit persistent passivity suppressing effects in the futility-induced passivity assay. (P-U) Effect on passivity in the futility-induced passivity assay after treatment with compounds shown in (I) compared to their same clutch, vehicle-treated controls. All error bars represent s.e.m. Two-tailed t-test with p-value indicated in each plot and number of fish above each bar. Numbers above bars denote the number of fish per group. All error bars denote s.e.m. *n.s.* p > 0.05, * p < 0.05, ** p < 0.01, *** p < 0.005, **** p < 0.001. Statistics for (J-O): One way ANOVA with Tukey’s multiple comparison test (vs.control). (J), N = 17 fish (control), 10 fish (ketamine), 8 fish (MK-801), p = 0.0086 (ketamine), 0.0115 (MK8-801). (K) N = 24 fish (control), 8 (fluoxetine), 8 (citalopram), 8 desipramine, p = 0.4790 (fluoxetine), 0.9990 (citalopram), 0.0036 (desipramine). (L) 16 fish (control), 8 (DOI), 7 (TCB-2), p = <0.0001 (DOI), 0.0008 (TCB-2). (M) N = 8 fish (control), 8 (ketamine), 8 (MK-801), p = 0.0103 (ketamine), <0.0001 (MK-801). (N) N = 8 fish (control), 8 (fluoxetine), 8 (desipramine), p = 0.2392 (fluoxetine), 0.0017 (desipramine). N = 16 fish (control), 8 (DOI), 8 (TCB-2), p = 0.8096 (DOI), 0.8295 (TCB-2).

**Figure S2.**
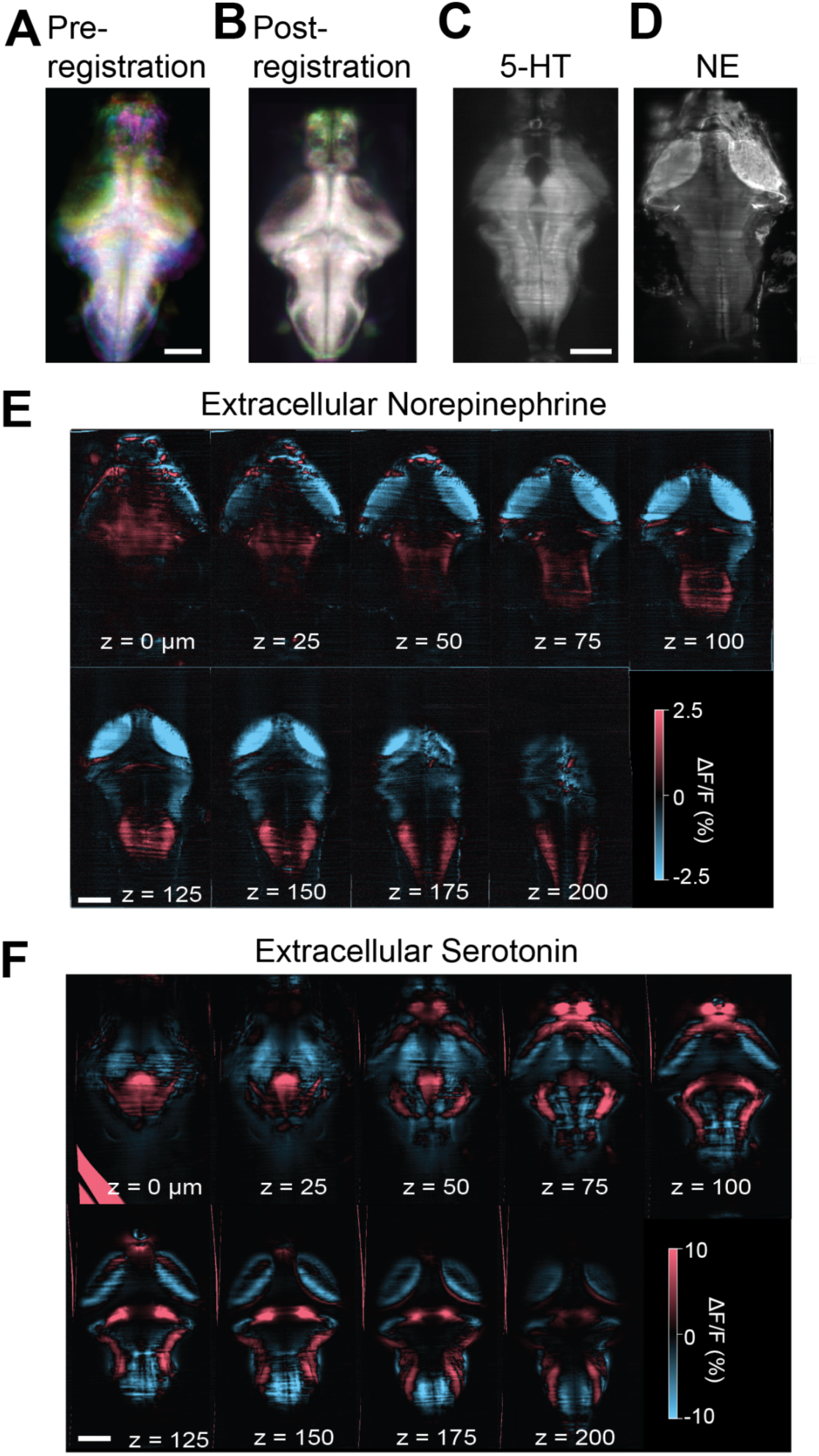
Relates to Fig. 2. (A-B) Max projections of all the fish used in Figure 2 before (A) and after (B) registration. (C-D) Anatomical stacks of (C) GRAB_5-HT_ and (D) GRAB-NE fish. (E) Average ΔF/F GRAB-NE changes across different brain planes after ketamine treatment (N = 2 fish). Scale bar 100μm. (F) Average GRAB_5-HT_ changes across different brain planes after ketamine treatment (N = 4 fish). Bright red regions outside of brain are registration artifacts. Scale bar 100μm.

**Figure S3.**
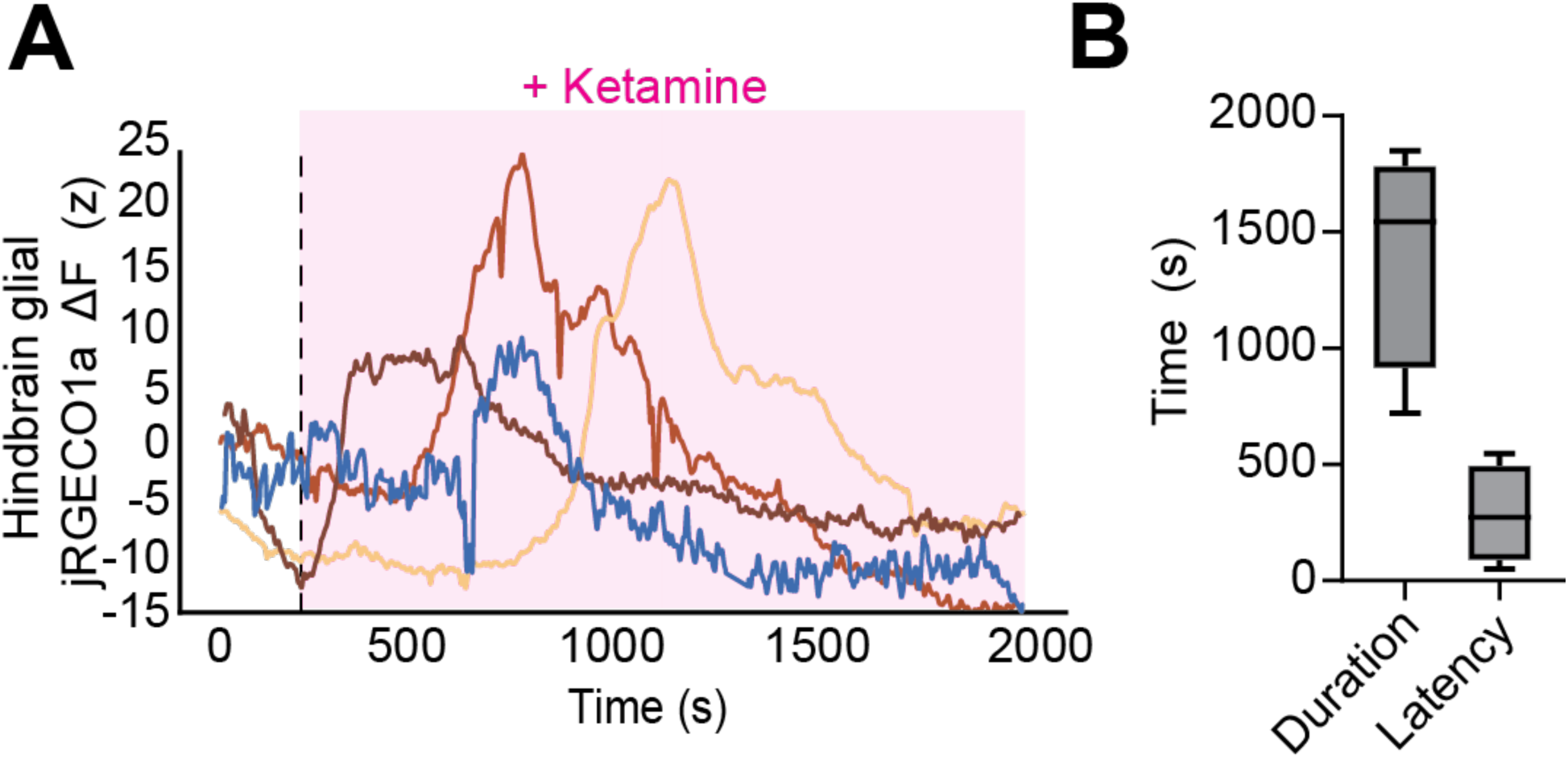
Relates to Fig. 3. (A) Mean astroglial fluorescence from hindbrain ROI for jRGECO in different fish after ketamine treatment. (B) Amplitude and latency (time to peak after ketamine administration) of the astrocytic calcium wave induced by ketamine (N = 4 fish). Mean ± SEM Amplitude = 1414s ± 242.9. Latency = 285.4 ± 105.7 s.

**Figure S4.**
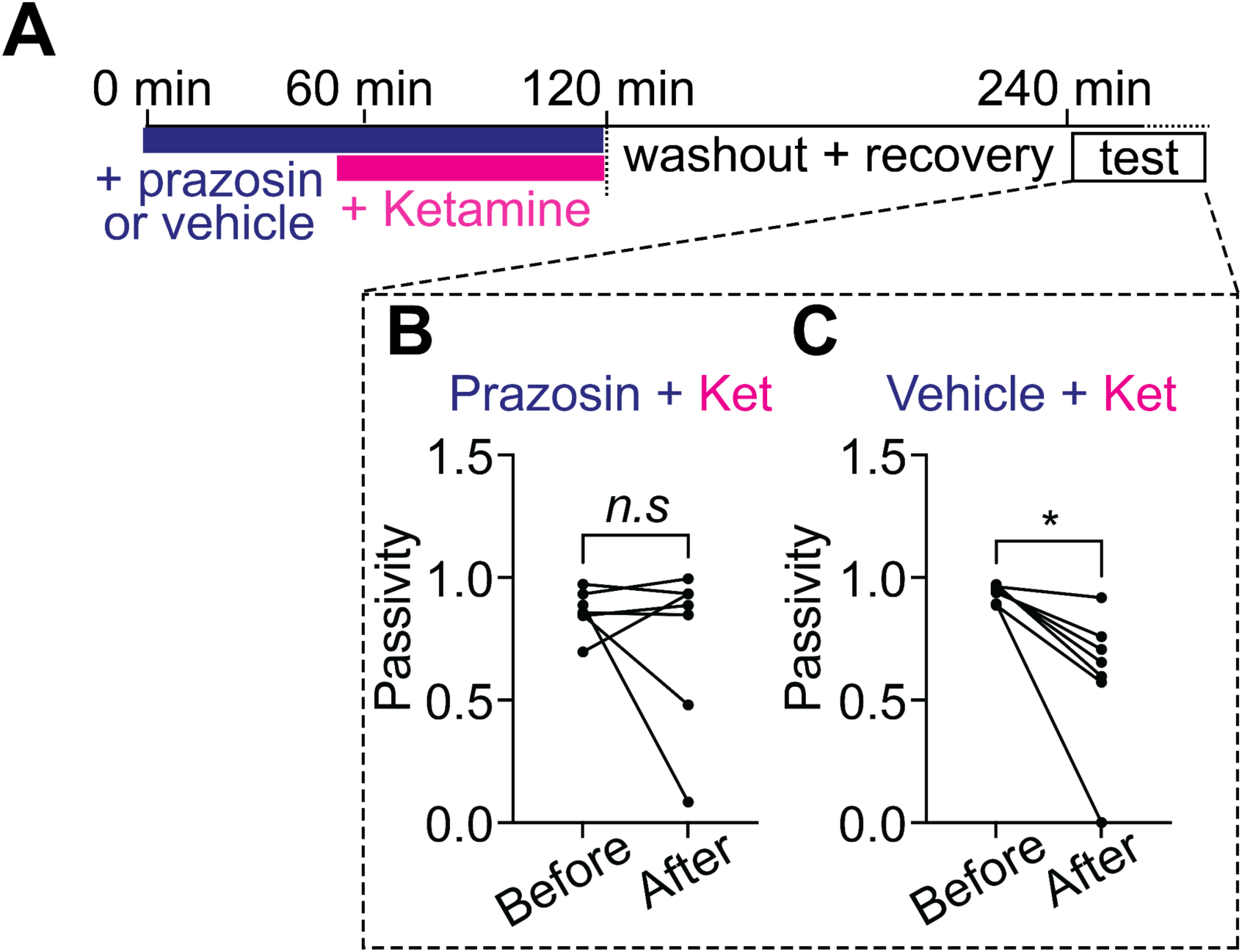
Relates to Fig. 4. (A) Schematic of experiments testing whether ketamine’s effects on astroglial Ca^2+^ and futility-induced passivity behavior depend on NE signaling. (B) Open-loop passivity in our assay for fish treated with ketamine and α1 adrenergic blocker prazosin (100 μM), before and after treatment. Incubation with prazosin blocks the effect of ketamine in our assay. Two-tailed paired t-test. N = 7 fish pairs. p = 0.3815. (C) Open-loop passivity in our assay for fish treated with ketamine and vehicle control, before and after treatment, as a clutch control for this experiment. Two-tailed paired t-test. N = 7 fish pairs. p = 0.0162.

**Figure S5.**
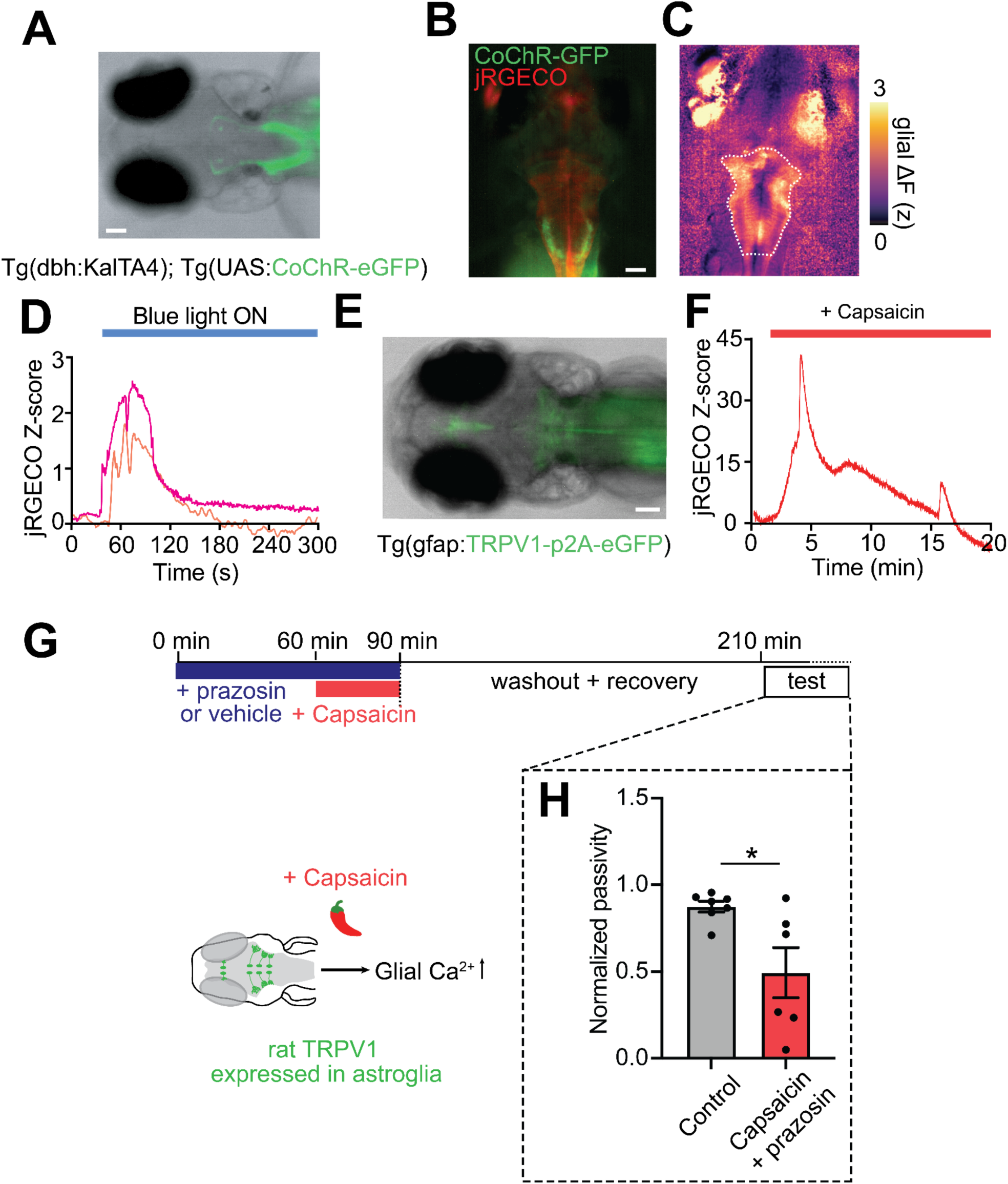
Relates to Fig. 5. (A) Overlaid fluorescence and transmitted light image of a larval zebrafish expressing eGFP-fused CoChR (a channelrhodopsin) under the *dbh* promoter. Expression in noradrenergic neurons evidence from fluorescence in anatomically distinct axon tracts. (B) Overlaid red-and green-channel fluorescence images showing double transgenic fish lines used for simultaneous imaging of glial cytosolic calcium and stimulating NE neurons. Fish express eGFP-fused CoChR under the *dbh* promoter and the red calcium indicator jRGECO1a under the *gfap* promoter in *Tg(dbh:KalTA4; UAS:CoChR-eGFP; gfap:jRGECO1a)*. (C) Imaging frame taken during blue-light (488 nm) stimulation of NE neurons while imaging astroglial cytosolic Ca. (D) Time-course of astroglial jRGECO1a signal in two different fish during blue-light stimulations. (E) Overlaid fluorescence and transmitted light image of a larval zebrafish expressing GFP-fused rat TRPV1 in astroglia under the *gfap* promoter (fish line *Tg(gfap:TRPV1-T2A-eGFP)*). Scale bars 50 μm. (F) Time-course of astroglial jRGECO1a signal in a fish expressing gfap:TRPV1 during treatment with capsaicin (2μM). Note the long-lasting elevation in astroglial calcium with similar duration to the acute effect of ketamine. (G) Timeline for experiments testing effects of stimulating glial calcium directly through chemogenetics in *Tg(gfap:TRPV1-T2A-eGFP)* fish while simultaneously blocking the alpha-1 adrenergic receptor with prazosin. Transgenic fish expressing rat TRPV1 in astroglia were treated with prazosin (100 μM) for 1h, followed by capsaicin treatment for 30 min. After 30 min, capsaicin was washed out and fish allowed to recover for 2 h before being tested in our assay. (H) Passivity normalized to untreated clutch controls. Prazosin treatment did not block the effect of capsaicin on futility-induced passivity. This suggests alpha-1 is only required upstream of glial calcium activation for the effects on futility-induced passivity. Mann-Whitney test. N = 7 fish (Control), 6 fish (Capsaicin [Cap] + Prazosin [Praz]). p = 0.0181.

**Figure S6.**
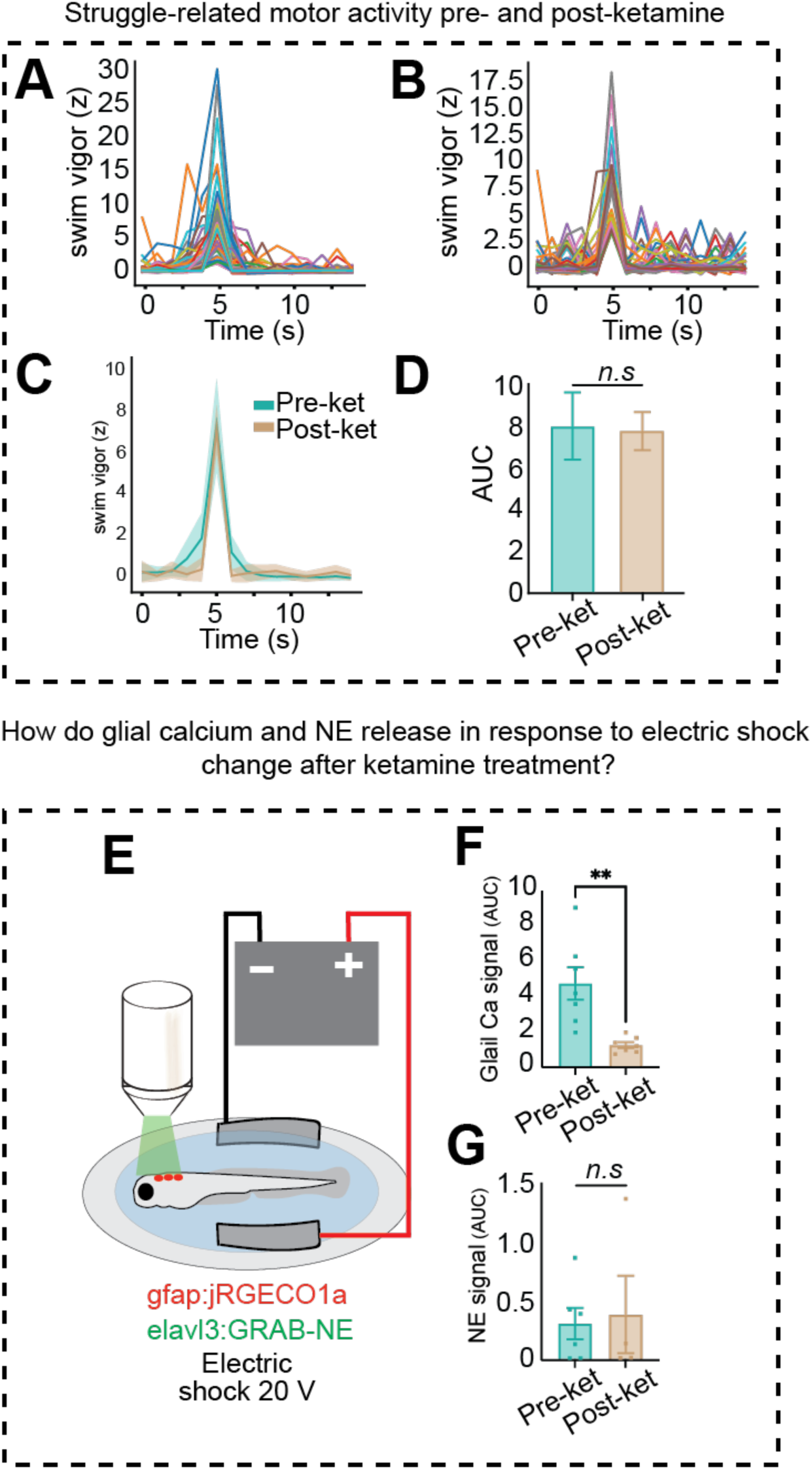
Relates to Fig. 6. (A) Set of struggles used in the struggle-triggered neuron and astroglia activity analysis from Figure 6B-E before ketamine (A) and after recovery from ketamine treatment (B). 60 struggles (pre), 56 struggles (post). (C) Average open-loop struggle swim traces for fish pre and post ketamine treatment. (D) Area under the curve (AUC) for the swim vigor traces shown in (C). Two-tailed t-test. N = 6 fish. p > 0.5. (E) Experimental setup to deliver mild electric shock. 20 V, 200ms stimuli were delivered while imaging either glial calcium with gfap:jRGECO1a or elavl3:GRAB-NE. (F) Area under the curve quantification for glial calcium ΔF/F traces in response to electric shock pre and post ketamine. Two-tailed t-test. N = 7 fish. p = 0.0020. (G) Area under the curve quantification for GRAB-NE ΔF/F traces in response to electric shock pre and post ketamine, suggesting a separate, unaffected source of NE during shock than during behavioral futility. Two-tailed t-test. N = 6 fish (pre), N = 4 fish (post). p = 0.8083.

**Figure S7.**
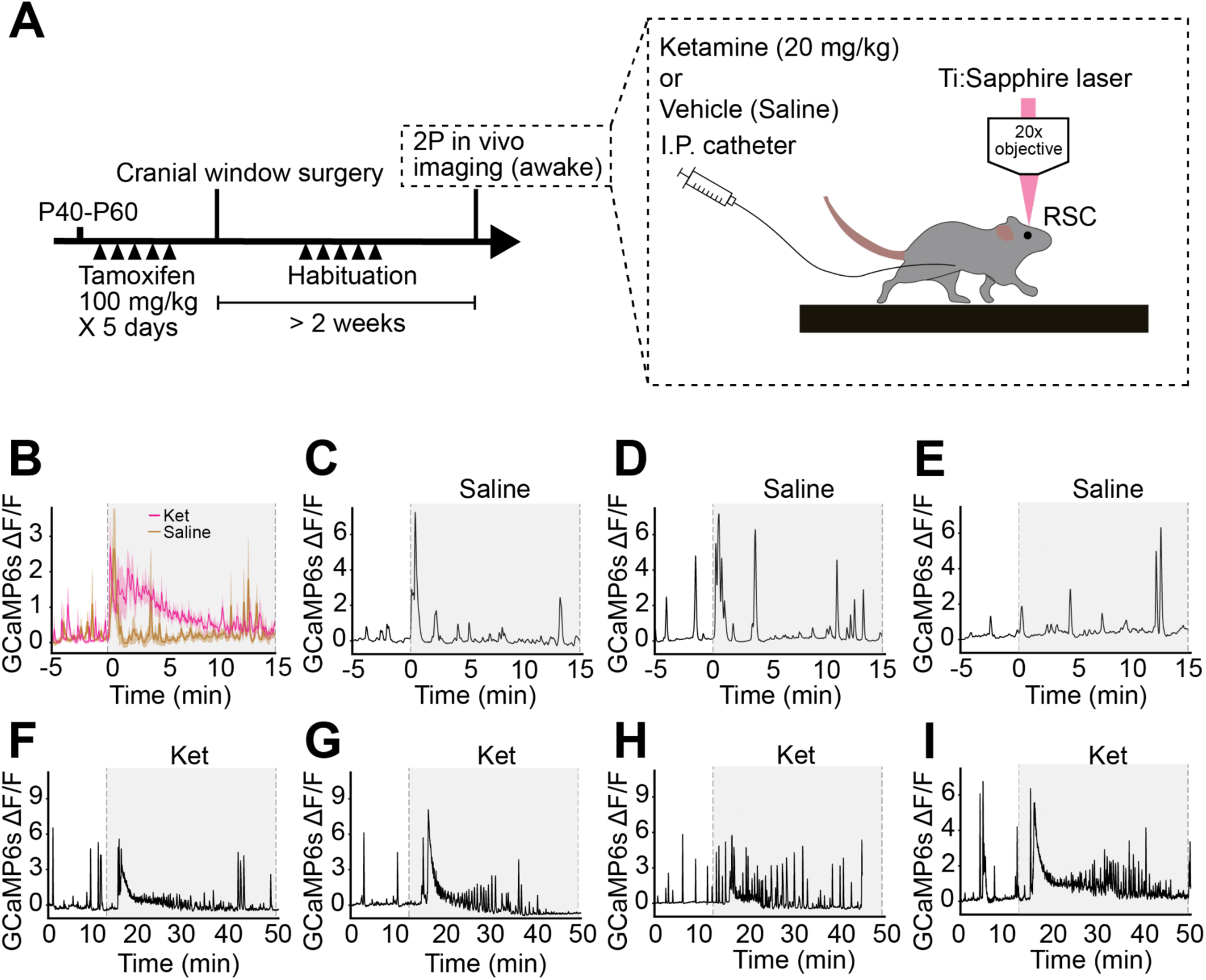
Relates to Fig. 7. (A) Schematic showing timeline of *in vivo* experiments (left) along with the setup to perform imaging of cortical astrocytes during I.P catheter injection in awake mice (right). (B) Average RSC GCaMP6f fluorescence trace for mice injected with control saline or ketamine at 15 min. (C-E) Individual traces corresponding to the average fluorescent trace in (B). (F-I) Individual traces corresponding to the average fluorescent trace in Figure 7C.

### Supplementary Videos

**Video S1. Example video of a control 7 dpf larval zebrafish in our assay for futility-induced passivity**. Example 10 s in rest (no forward drift, feedback), followed by 10s in closed loop (forward drift, feedback) and 10 s in open loop (forward drift, no feedback). The 3 intervals are discontinuous and have been merged for visualization purposes.

**Video S2. Example video of a 7 dpf larval ketamine-treated larval zebrafish in our assay for futility-induced passivity following recovery from the acute effects of ketamine treatment.** Example 10s in rest (no forward drift, feedback), followed by 10 s in closed loop (forward drift, feedback) and 10 s in open loop (forward drift, no feedback). The 3 intervals are discontinuous and have been merged for visualization purposes.

**Video S3. Example video of an astroglial calcium wave in response to acute ketamine treatment.** ΔF of whole-brain jRGECO1a signal expressed under the GFAP promoter imaged with an epifluorescence scope at 2 Hz. Ketamine was introduced to the water bath after 5 minutes. Scale bar 50 μm.

**Video S4. Example video of a cortical astrocytic calcium wave in response to acute ketamine treatment in awake mice.** ΔF of GCaMP6s signal expressed under the Aldh1l1 promoter imaged with a two-photon microscope through a cranial window in the mouse retrosplenial cortex. Ketamine 20 mg/kg was injected IP after 15 minutes. Scale bar 50 μm.

**Table S1.**
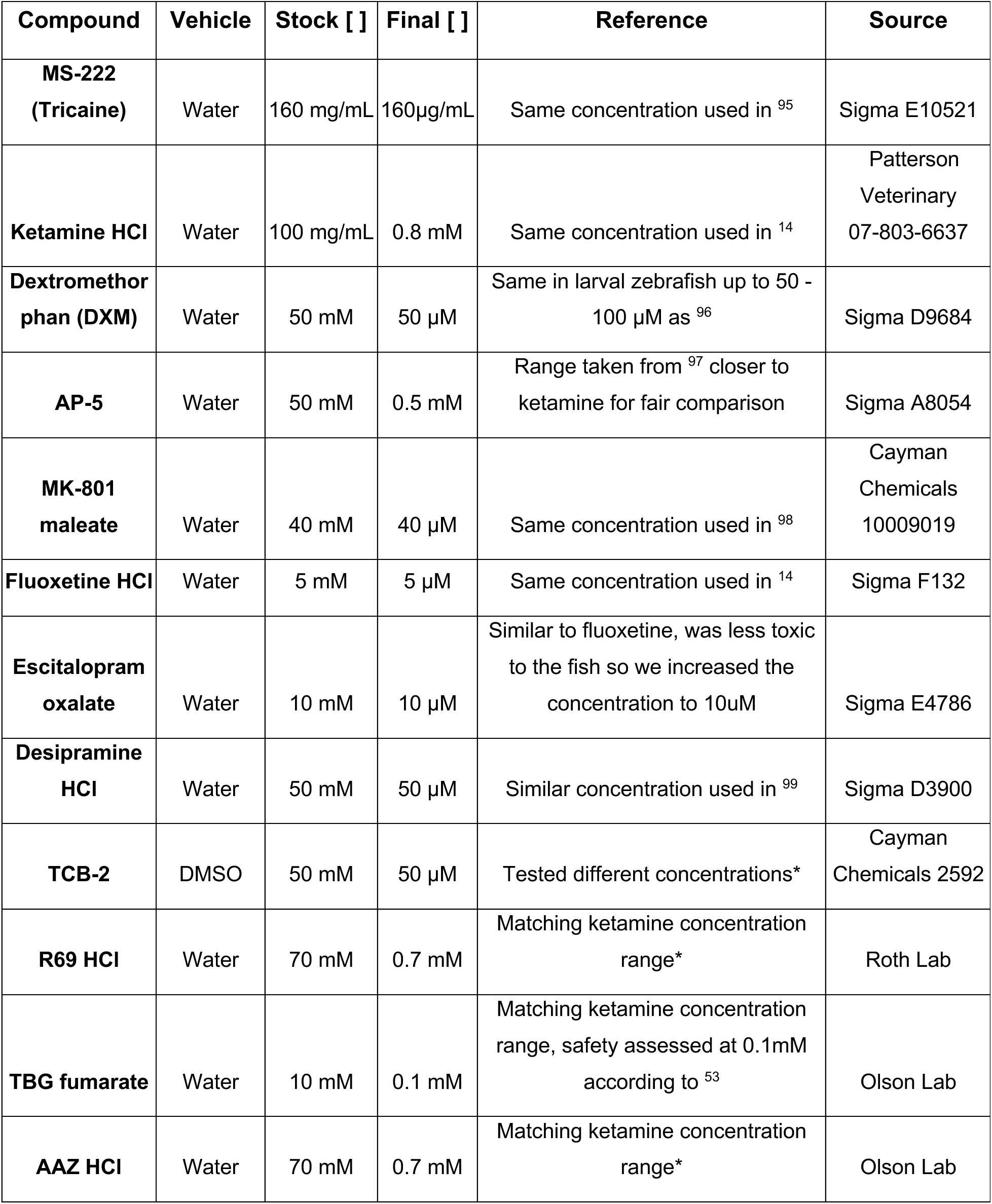
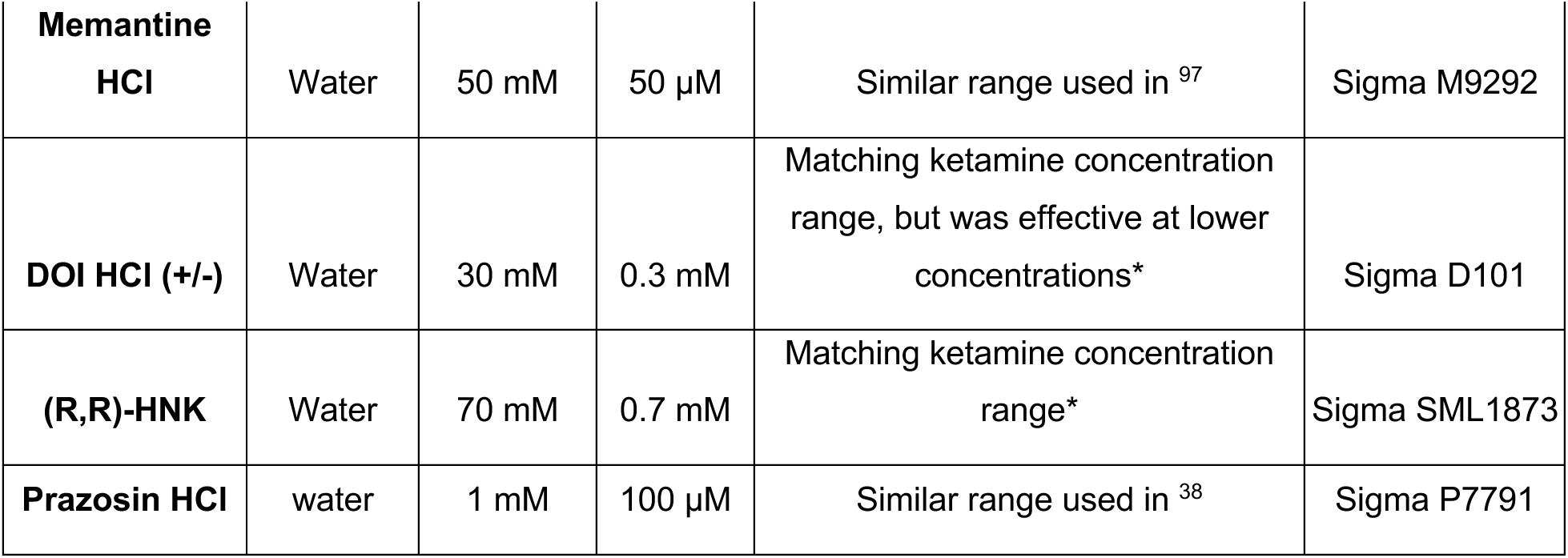
List, sources, and concentrations of compounds used in pharmacological experiments. * denotes no record of testing in larval zebrafish.

## Notes

### Summary of Updates

Updated title, authors affiliations updated, abstract updated, updated text, supplemental files updated, main figures updated, updated text

